# Chemical-genetic interrogation of RNA polymerase mutants reveals structure-function relationships and physiological tradeoffs

**DOI:** 10.1101/2020.06.16.155770

**Authors:** Anthony L. Shiver, Hendrik Osadnik, Jason M. Peters, Rachel A. Mooney, Peter I. Wu, James C. Hu, Robert Landick, Kerwyn Casey Huang, Carol A. Gross

**Author notes:** Lead contact: Carol Gross.

## Abstract

The multi-subunit bacterial RNA polymerase (RNAP) and its associated regulators carry out transcription and integrate myriad regulatory signals. Numerous studies have interrogated the inner workings of RNAP, and mutations in genes encoding RNAP drive adaptation of *Escherichia coli* to many health- and industry-relevant environments, yet a paucity of systematic analyses has hampered our understanding of the fitness benefits and trade-offs from altering RNAP function. Here, we conduct a chemical-genetic analysis of a library of RNAP mutants. We discover phenotypes for non-essential insertions, show that clustering mutant phenotypes increases their predictive power for drawing functional inferences, and illuminate a connection between transcription and cell division. Our findings demonstrate that RNAP chemical-genetic interactions provide a general platform for interrogating structure-function relationships *in vivo* and for identifying physiological trade-offs of mutations, including those relevant for disease and biotechnology. This strategy should have broad utility for illuminating the role of other important protein complexes.

## Introduction

Multi-subunit RNA polymerases are responsible for transcription in all organisms. The core RNA polymerase (RNAP) enzyme (β’, β, α_2_, ω) is conserved across all domains of life (Jokerst et al., 1989; Lane and Darst, 2010; Sweetser et al., 1987). Bacterial-specific initiation factors, called sigmas (σs), transiently associate with the core complex to provide promoter recognition and assist in melting promoter DNA during initiation (Gruber and Gross, 2003). During elongation, RNAP associates with NusA, which enhances pausing and intrinsic termination at specific sequences (Artsimovitch and Landick, 2000), and NusG (Spt5 in archaea and eukaryotes), the only universally conserved elongation factor, which modulates elongation and ρ-dependent termination (Burova et al., 1995; Li et al., 1993). Termination in eubacteria is facilitated either by RNA structure (intrinsic termination) or by the termination factor ρ, which uses its helicase activity to release the transcript and recycle the RNAP complex (**Figure 1A**). Additionally, bacterial RNAPs differ from archaeal and eukaryotic RNAPs, for which the core enzymes acquired peripheral subunits (e.g., Rpb4,5,7-10,12 in RNAPII), by instead having acquired lineage-specific insertions in β’ and β (called sequence insertions 1-3 (SI1-3) in *E. coli*) whose functions remain largely unknown (Artsimovitch et al., 2003; Lane and Darst, 2010).

**Figure 1:**
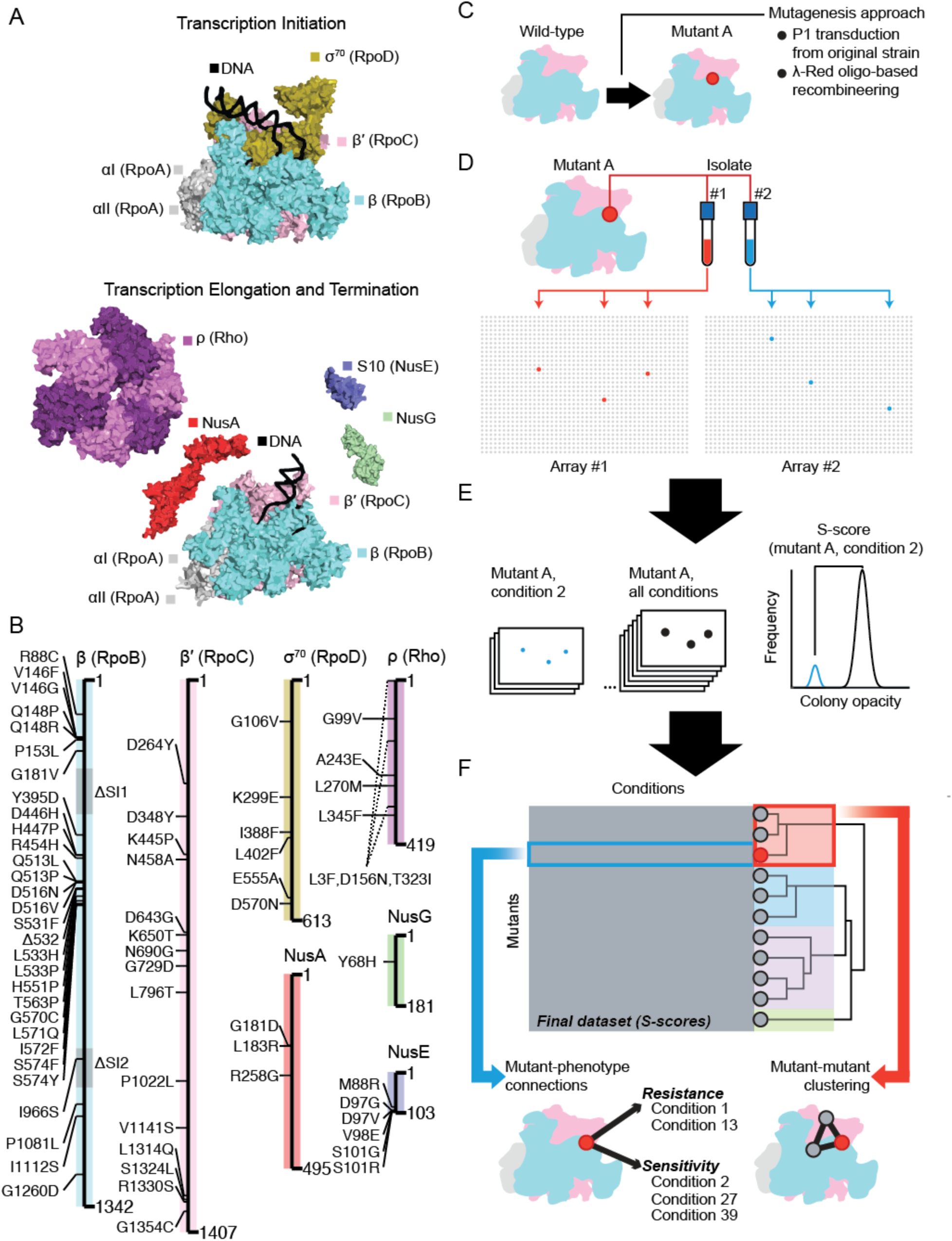
A chemical-genetic screen focused on the bacterial transcription machinery dissects structure-function relationships of RNA polymerase *in vivo*. A) The core essential subunits of RNA polymerase (α_2_ββ′) form a complex with sigma factors such as σ^70^ during transcription initiation. During elongation, factors such as NusA, NusG, NusE, and ρ associate with RNAP to regulate pausing and termination. B) A library of 68 mutations in RNAP was created in an isogenic background to dissect RNA polymerase function *in vivo*. C) Mutations were introduced *de novo* using oligo-based recombineering or transduced from the original isolate using P1*vir*. D) Independent isolates of the same mutation (red and blue) were split between parallel sets of colony arrays (Array #1 and Array #2). Each isolate was arrayed in triplicate and the position of each mutant was randomized between the two arrays. The randomized positions of the biological replicates are shown as red colonies in a 32 row × 48 column array of colonies. The remaining mutants are gray. E) At an appropriate time for each condition, a single image was taken of every plate and colony opacity was estimated using image analysis software. Following appropriate normalizations and filtering steps, the distribution of colony opacity measurements for a given mutant/condition pair were compared to the entire distribution of opacity measurements of the mutant across all conditions to generate an S-score. The S-score is a modified t-statistic that measures the significance of the difference between colony opacity (closely related to colony size) of a specific mutant/condition pair (blue colonies, blue distribution of colony opacity) and the control of the same mutant across all conditions (black colonies, black distribution of colony opacity). In this example, colony opacity is lower on the given condition, leading to negative S-scores that are interpreted as chemical sensitivity. Conversely, higher colony opacities would lead to positive S-scores that are interpreted as resistance. Importantly, S-scores are proportional to the statistical significance of an interaction, not the direct magnitude of the interaction itself. F) The final dataset was a 68 × 83 matrix of mutant × condition S-scores. Individual S-scores were investigated to identify new mutant-phenotype connections and hierarchical clustering of mutants was used to assign new functions to mutations.

The central role played by this enzyme complex, both in orchestrating transcription and integrating diverse signals, is reflected in the pleiotropic phenotypes that arise from mutations in RNAP. Efforts to evolve *E. coli* in maladapted environments, such as growth on glycerol (Cheng et al., 2014), ethanol (Haft et al., 2014), or at elevated temperatures (Tenaillon et al., 2012), have all recovered mutations in RNAP as a predominant class, highlighting the intimate ties of RNAP to a wide range of cellular processes. Adaptation in these conditions is highly relevant for biotechnology applications, as directed mutagenesis of RNAP could serve as a rapid means of adapting bacteria to new production environments (Alper and Stephanopoulos, 2007). However, without a deeper understanding of how RNAP mutations affect cellular physiology, it will be difficult to rationally design mutations and hard to predict whether mutations will have unintended physiological side effects.

Chemical-genetic screens measure the effect of stressful environments, such as the presence of an antibiotic, on growth across a large library of mutations (Brochado and Typas, 2013). By discovering novel growth phenotypes and identifying mutants with highly correlated growth phenotypes across conditions, such screens generate new hypotheses regarding biological pathways and gene functions (Nichols et al., 2011). Chemical screens can also be used to analyze a large collection of mutations in a single protein complex (Braberg et al., 2013), wherein discovery of new phenotypes and correlations between phenotypic profiles make possible *in vivo* structure-function analyses based on the effects of mutations on cellular physiology. By interrogating chemical-genetic interactions across a wide range of environments, these screens are also uniquely situated to identify the secondary effects of adaptive mutations.

In this work, we conducted a chemical-genetic screen focused on RNAP mutations in *E. coli* K-12, with the goal of interrogating connections between RNAP and cellular physiology and dissecting *in vivo* structure-function relationships within RNAP and its associated factors. We generated an isogenic library of 68 unique mutations in RNAP and essential transcription factors and screened the library in 83 unique conditions to generate a chemical-genetic dataset that we integrated with existing data from the Keio library of all nonessential gene deletions. We confirmed that mutations in RNAP are highly pleiotropic, with altered sensitivities to antibiotics that target peptidoglycan synthesis, folate biosynthesis, DNA replication, and translation. We shed light on the effect of understudied features of RNAP like β-SI2 on transcription. Finally, we identified a cross-resistance phenotype of RNAP mutations based on stabilization of FtsZ. Taken together, these data illustrate the power of chemical-genetic screens to illuminate *in vivo* structure-function landscapes.

## Results

### Construction of a library of strains with chromosomal mutations in the transcription machinery

Decades-long study of the *E. coli* transcriptional apparatus has generated a large set of mutations with diverse phenotypes, particularly in the two largest subunits of RNAP—β’ and β—and to a lesser extent in σ^70^, NusA, NusG, and ρ. Unfortunately, the phenotypes of these mutant strains are not immediately comparable, as RNAP mutations are in diverse genotype backgrounds and are often only found as episomal merodiploids with a wild-type chromosomal copy. Following a literature review to manually annotate and collate the existing mutants, we chose and successfully reconstructed 68 mutations (**Figure 1B, Supplemental Table 1**) at their endogenous locus in the BW25113 strain background, enabling comparison with published chemical genomics datasets (Nichols et al., 2011; Shiver et al., 2016). Some mutations were introduced via transduction using a closely linked antibiotic resistance cassette; others were reconstructed by λ-Red oligo-mediated recombineering into a strain containing that cassette (**Figure 1C**). Rif^R^ mutants that confer resistance to rifampicin, and M^+^ mutants that restore growth in minimal media to strains that lack or are deficient in the mediators of the stringent response ppGpp and DksA (Murphy and Cashel, 2003), are overrepresented in this collection (32/68) because they could be identified by selection. Some M^+^ mutants have been shown to form innately unstable open promoter RNAP complexes *in vitro*, mimicking the effects of ppGpp and DksA binding to RNAP, likely explaining their phenotype in minimal media (Rutherford et al., 2009). Our library also included a wide variety of other mutants that ensured our capacity to detect diverse phenotypic profiles.

### A chemical-genetic screen of the transcription library reveals residue-level phenotypes of transcription mutants *in vivo*

We performed a chemical-genetic screen of the arrayed mutant library using sub-inhibitory concentrations of 83 chemical stressors that overlapped with previous screens (Nichols et al., 2011; Shiver et al., 2016). The screen was performed in duplicate with technical and biological replicates of the mutant strains (**Figure 1D**) as well as control strains with the antibiotic marker alone and a subset of deletion strains from the Keio library. We quantified colony opacity from images of the colony arrays at a single time point to calculate S-scores, a measure of sensitivity or resistance in each condition (**Figure 1E**) (Collins et al., 2006; Kritikos et al., 2017). S-scores were internally reproducible (*r*=0.73 for the same mutant compared across screens), and S-scores for the nonessential gene deletions were correlated with those determined in a previous screen (*r*=0.65) (Shiver et al., 2016). This final S-score dataset was used in subsequent analyses examining chemical sensitivities and mutant-mutant correlations (**Figure 1F**). The entire dataset is available in an interactive, searchable format on the Ontology of Microbial Phenotypes website https://microbialphenotypes.org/wiki/index.php?title=Special:ShiverSpecialpage.

We used a cutoff based on hierarchical clustering of the S-scores to define 23 statistically significant clusters of transcription mutants (**Supplemental Figure 1**). Mutations largely clustered with others in the same gene (**Figure 2A**), except for mutations in β and β′, which frequently clustered together. β and β′ are interwoven to form the core of the RNAP complex, and many of the mutations in these subunits are found on either side of the same DNA binding cleft. Clustering of mutations in these subunits likely reflects their tight functional coordination in the complex. Setting aside interactions between β and β′, only one out of 171 co-clustering interactions was between mutations in different subunits (odds ratio=205, *p*=10^-53^). This interaction was between β-I1112S and NusA-R258G, which comprised cluster 22 (**Figure 2A,B**). β-I1112S and NusA-R258G were isolated from the same screen for ethanol tolerance (Haft et al., 2014), and their co-clustering confirmed that shared effects on the transcription cycle can drive clustering of mutations in the chemical genomic dataset beyond occurring in the same subunit.

**Figure 2:**
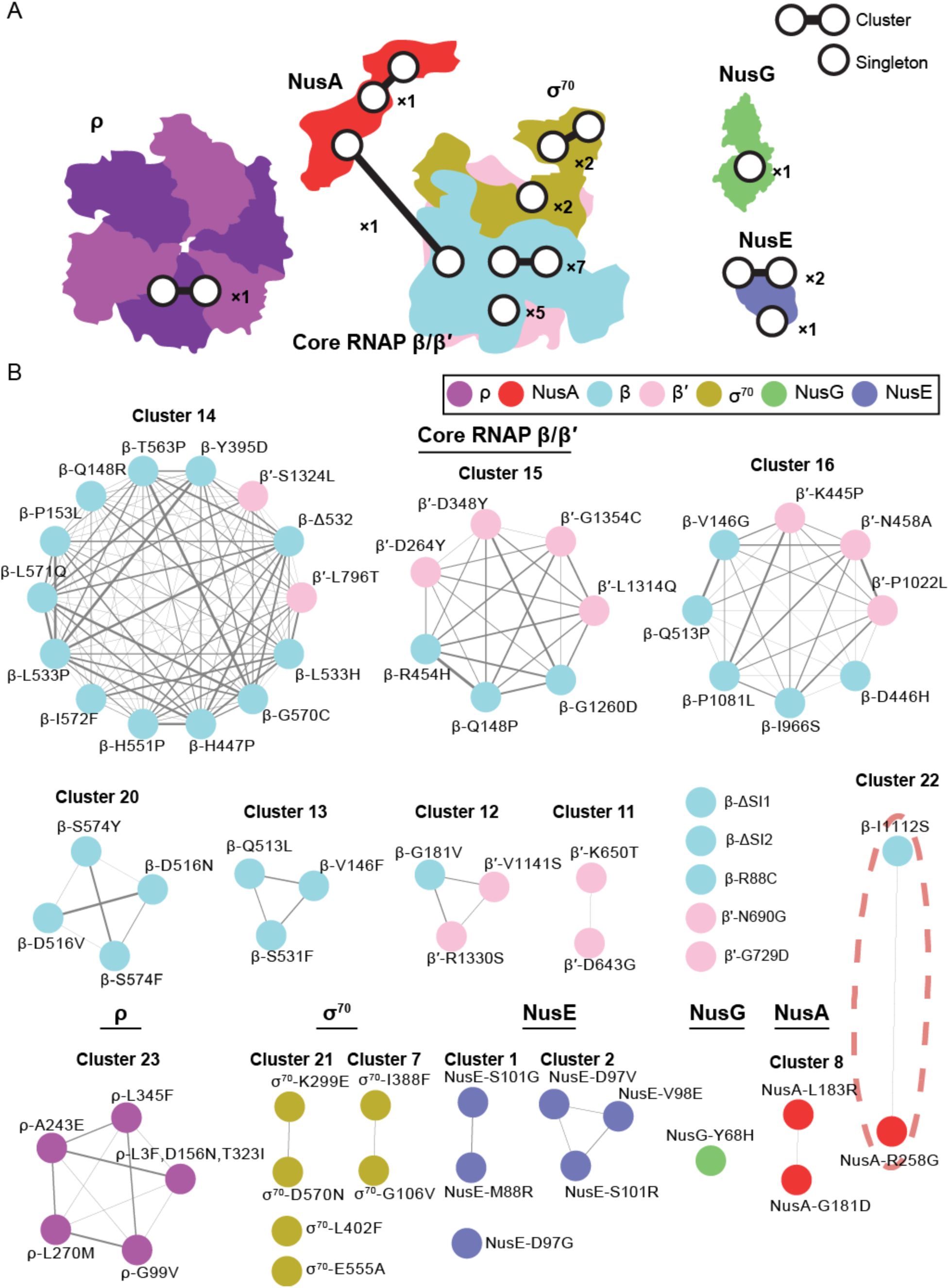
Mutations in the transcription machinery cluster mainly according to subunits. A) Summary statistics of clustering the dataset are superimposed onto the structures of the 7 RNAP proteins with mutations included in the chemical-genetic screen. Two points connected by a line represent a cluster of ≥2 mutations as defined by the screen. A single point represents a singleton mutation with no significant correlation with any other transcription mutant. Mutations in ρ form a single cluster. Most NusA mutations form one intra-subunit cluster, but one NusA mutation clusters with a mutation from core RNAP. Mutations in σ^70^ form two clusters and two are singletons. Mutations in β/β′ form seven clusters and 5 mutations are singletons. The lone mutation in NusG is a singleton. Mutations in NusE form two clusters and one is a singleton. B) The full clusters are shown color-coded and grouped according to subunit. The width of the edge connecting two mutations is proportional to their correlation in the dataset (Pearson’s *r*). The sole cluster that connects mutations from different subunits (cluster 22) is highlighted in red.

We calculated the enrichment of chemical-genetic interactions among clusters (**Supplemental Table 2**) and mutant classes (**Supplemental Table 3**). Focusing on the three largest clusters of β and β’ mutations (out of 7 total) (**Figure 3A)**, we found that each cluster could be associated with unique chemical-genetic interactions made by the mutants. The largest cluster (14) was enriched for sensitivities to aminoglycosides such as spectinomycin (**Figure 3B**). Cluster 15 was enriched for resistance to the tetracycline family of antibiotics (**Figure 3C**). Cluster 16 was enriched for sensitivity to tetracycline (**Figure 3C**) and enriched for resistance to trimethoprim (**Figure 3D**). Interestingly, this clustering did not necessarily follow the transcriptional classifications of the mutations: cluster 14 is comprised of both Rif^R^ and M^+^ mutations on the β-side of the RNAP cleft, cluster 15 is comprised predominantly of M^+^ mutations on the β’-side of the RNAP cleft, and cluster 16 contains mutations spread across the complex and with different known phenotypes (**Figure 3A**). Our observation of mutant clusters that are not aligned with previously defined classifications suggests that the chemical-genetic interactions in our dataset contain more detailed information regarding the effects of these diverse mutations on cellular physiology, a proposition we explore in the following sections as we investigate the phenotypes of specific RNAP mutations.

**Figure 3:**
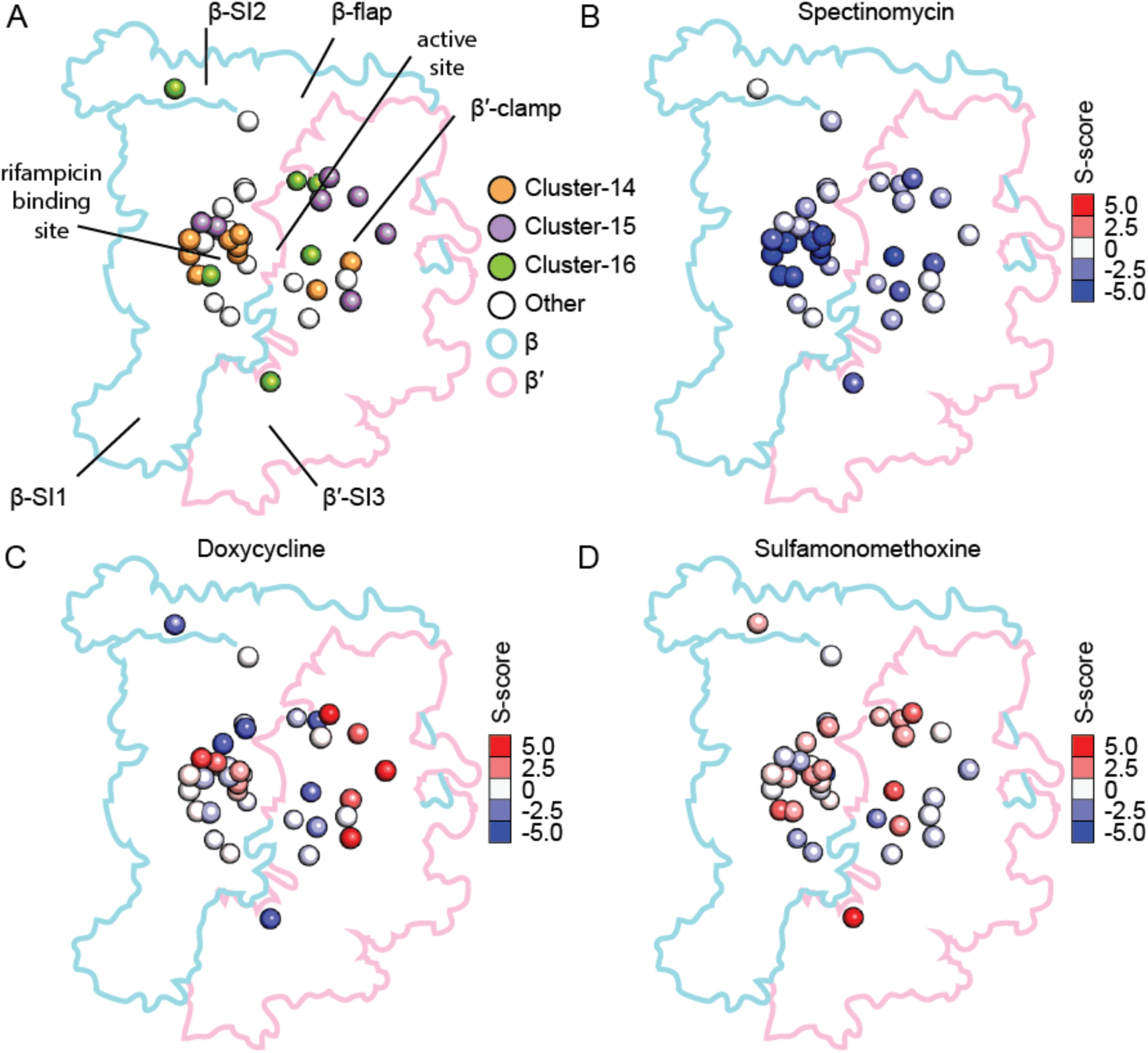
RNA polymerase mutants in the same subunit can be dissected into clusters based on chemical sensitivities. A) Three major clusters of point mutations within the core complex of β and β′ were identified. Cluster 14 (orange) is centered around the rifampicin binding site, cluster 15 (purple) is found mostly in the β′-clamp, and cluster 16 (green) is distributed throughout the complex, including β′-SI3, the active site, and β-SI2. Point mutations in RNAP not included in clusters 14-16 are shown in white. The alpha carbon of mutated residues is shown as a sphere in the structure. B) Sensitivity to spectinomycin is enriched in cluster 14. The alpha carbons of mutated residues are shown as spheres colored by the magnitude and direction of the mutant’s S-score in the dataset. Red indicates resistance and blue indicates sensitivity. C) Resistance to doxycycline is enriched in cluster 15 and sensitivity to doxycycline is enriched cluster 16. Mutations are presented in the structure as in (B) D) Resistance to sulfamonomethoxine is enriched in cluster 16. Mutations are presented as in (B).

### Chemical-genetic profiling the β subunit non-essential sequence insertions reveals environmental sensitivities

The chemical-genetic phenotypes of deletions of the large, non-essential sequence insertions, β-SI1 and β-SI2, were outliers among the transcription mutants (**Figure 2B**), suggesting that their impacts on RNAP function are unique within the library. β-ΔSI1 correlates with auxotrophic gene deletions, consistent with a role in the binding and function of the transcription factor DksA (Parshin et al., 2015). By contrast, β-ΔSI2 was not significantly correlated with any of the other transcription mutants or any mutants from the larger gene deletion library, making it difficult to ascertain its function by comparison to well-characterized mutants.

Although β-SI2 has been proposed to be dispensable for RNAP function (Borukhov et al., 1991; Nene and Glass, 1984; Severinov et al., 1992), we identified significantly negative S-scores for β-ΔSI2 in multiple treatments, including ethanol, trimethoprim, and hydroxyurea. To explore these potential sensitivities further, we monitored growth of β-ΔSI2 and its parental control in LB medium with increasing concentrations of all three compounds. At sub-lethal doses of ethanol, growth of the parental control slowed near the transition to stationary phase (**Figure 4A**). For β-ΔSI2, this phenotype was more pronounced and occurred at lower concentrations (**Figure 4A**). We found a similar chemical-genetic interaction with trimethoprim and hydroxyurea. In each case, growth of β-ΔSI2 was impacted by lower concentrations of the compounds but the mutant had no discernible effect on the MIC (**Figure 4B**). Thus, our screen revealed conditions that highlight the importance of β-SI2 to RNAP function.

**Figure 4:**
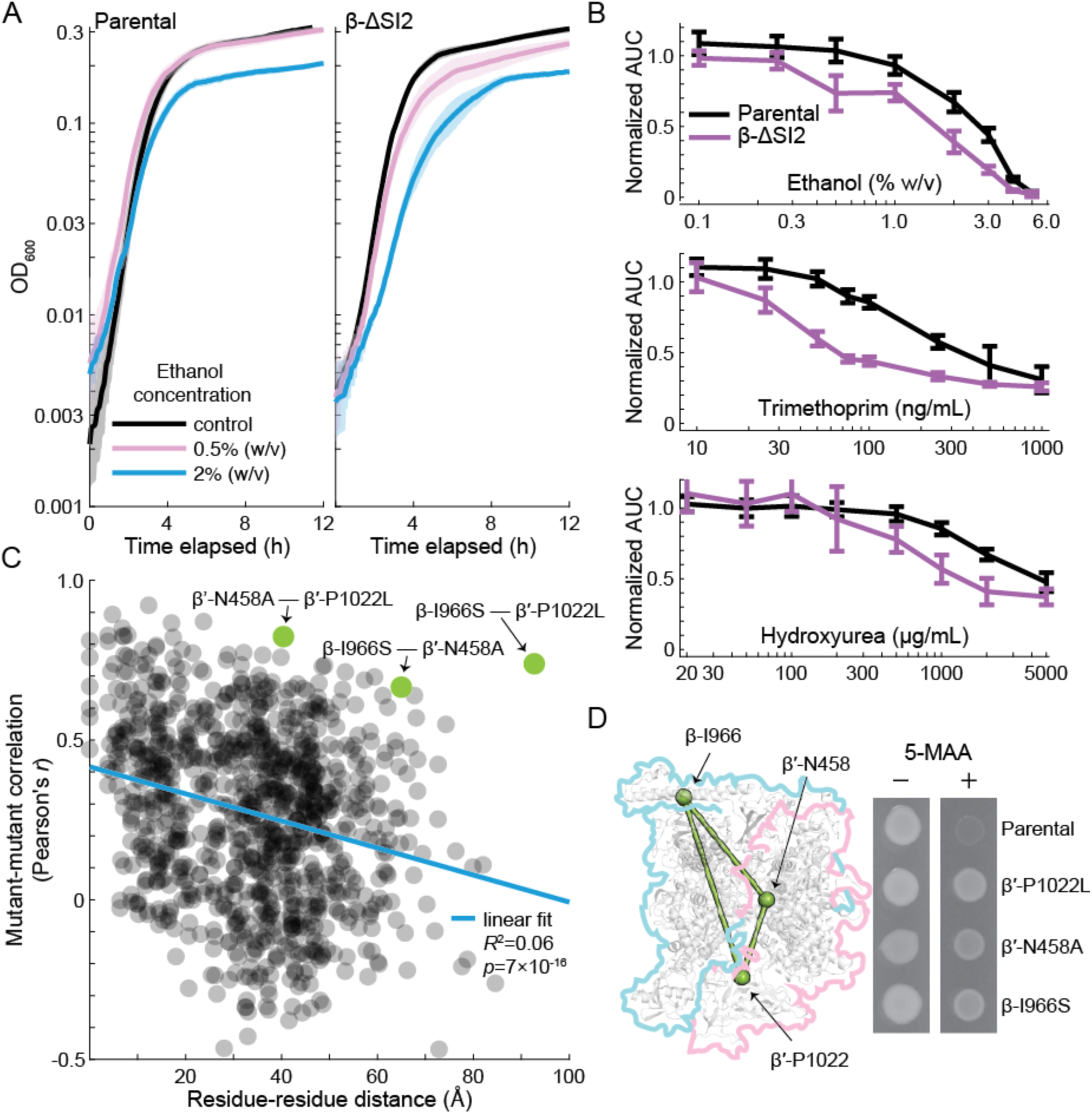
Mutations in β-SI2 have distinct phenotypes that include hyper-attenuation at the *trp* locus. A) β-ΔSI2 has mild sensitivity to ethanol. Left: sublethal doses of ethanol hampered growth of the parental *E. coli* strain starting during the transition to stationary phase. Right: β-ΔSI2 has a more pronounced response to ethanol that begins at a lower concentration of ethanol. B) β-ΔSI2 has mild sensitivity to ethanol, trimethoprim, and hydroxyurea. The normalized area under the curve (AUC) of growth curves as shown in (A) is plotted against the concentration of the compound. The AUC was calculated by integrating OD_600_ over time and normalizing by the AUC of the same strain without added drug. For ethanol (top), trimethoprim (middle), and hydroxyurea (bottom), growth of β-ΔSI2 was affected at lower (sub-inhibitory) concentrations, but the minimum inhibitory concentration remained the same. Error bars represent 95% confidence intervals. C) Mutant-mutant correlations show a weak association on distance between the residues in the RNA polymerase structure. Mutant-mutant correlations were calculated from Pearson’s *r* using the chemical-genetic dataset. Residue-residue distance was calculated as the 3D distance between alpha carbons of residues with mutations in the dataset. The PDB structure 4JKR was used as a basis for distance calculations. A three-mutant clique comprised of β-I966S, β′-N458A, and β′-P1022L was an exception to this rule, with high mutant-mutant correlations despite containing the largest inter-residue distance in the library (β-I966S to β′-P1022L). D) The correlations among β-I966S, β′-N458A, and β′-P1022L were predictive of a shared hyper-attenuation phenotype that was originally identified for β′-P1022L (Weilbaecher et al., 1994). In a Δ*trpR* background, expression of the *trp* locus is mainly controlled by attenuation. Hyper-attenuation reduces *trp* expression and makes cells resistant to a toxic analogue of a tryptophan biosynthesis intermediate, 5-methyl anthranilic acid (5-MAA).

### Phenotypic clustering uncovers residue-level RNAP phenotypes

Similar to a previous study examining the genetic interactions of yeast RNA Pol II mutations (Braberg et al., 2013), there was a statistically significant but weak negative correlation between distance in the structure and pairwise phenotypic correlations between mutants in our dataset (**Figure 4C**) (Pearson’s *r*=-0.25, *R*^2^=0.06, *p*=7×10^-16^). Excitingly, this correlation provided enough information to use pairwise comparisons of mutant phenotypic vectors as a distance constraint to improve structural modeling of the core complex (Braberg et al.).

Cluster 16 includes a high-correlation clique of three mutations: β-I966S, β′-N458A, and β′-P1022L. This clique was exceptional in that the phenotypic profiles of its members were highly correlated despite occurring in separated structural features of the RNAP complex (**Figure 4C**). Since β′-P1022L was isolated in a screen for increased transcription attenuation (Weilbaecher et al., 1994), we tested whether the other two mutants in the clique share this phenotype. As predicted by their high phenotypic correlation with β′-P1022L, both β-I966S and β′-N458A were resistant to 5-MAA (**Figure 4D**), indicative of increased transcription attenuation at the *trp* locus.

### The resistance of β-P153L to mecillinam and A22 is not due to a classical stringent response

Both cluster 14 and the M^+^ class of mutations were enriched for resistance to the cell wall-targeting drugs mecillinam and A22 (**Supplementary Tables 2,3**). At the sub-lethal doses used in our screen, A22 resistance was mostly restricted to a subset of cluster 14 near the rifampin binding pocket, while mecillinam resistance was found throughout the complex (**Figure 5A**). To investigate these connections mechanistically, we focused on β-P153L, an M^+^ mutant in cluster 14 with the highest positive S-score for A22 and robust resistance to mecillinam (**Figure 5A**).

**Figure 5:**
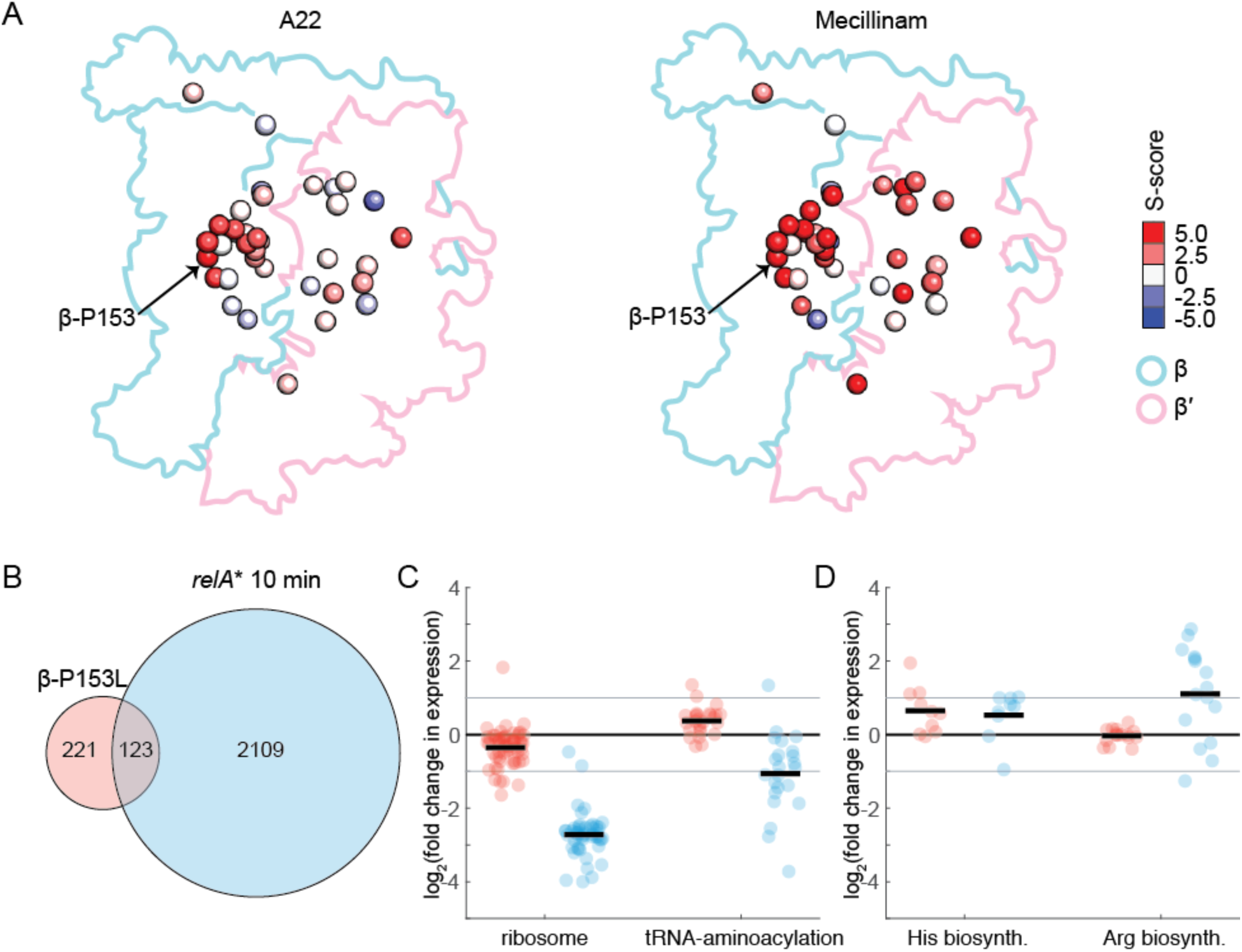
Gene expression in β-P153L only weakly overlaps with the stringent response. A) Resistance to mecillinam and A22 was enriched among M^+^ mutants and cluster 14. Left: In the screen, resistance to A22 was more concentrated in cluster 14 mutants around the rifampicin binding pocket. Right: at the sub-inhibitory concentration of mecillinam used in the screen, resistance was widespread throughout β and β′. The mutation β-P153L had the highest level of resistance to A22 and had high resistance to mecillinam. Alpha carbons of residues with mutations are colored according to their S-score in the chemical-genetic dataset. B) There was a small degree of overlap in the significantly differentially expressed genes in β-P153L (red, data collected in this study) and 10 minutes post-induction of a constitutively active *relA*\* allele (blue, from (Sanchez-Vazquez et al., 2019)). 36% of differential expressed genes in β-P153L overlapped with the much larger set from *relA\**. C) Repression of ribosome gene expression was weak in β-P153L (red), but consistent with results from *relA*\* induction (blue) and with the lower growth rate of the mutant in LB. The activation of tRNA-aminoacylation genes is opposite to the repression in the *relA*\* dataset. Individual genes are plotted as circles. The set average is shown as a sold black line. 2-fold changes in expression are marked with a solid gray line. D) Activation of genes involved in histidine biosynthesis in β-P153L (red) was consistent with the stringent response and comparable in magnitude to the *relA\** dataset (blue). However, histidine biosynthesis was the only biosynthetic pathway differentially expressed in the β-P153L mutant; for example, the lack of differential expression of the arginine biosynthetic pathways in β-P153L contrasted with the *relA\** dataset. Individual genes are plotted as circles. The set average is shown as a sold black line. 2-fold changes in expression are marked with a solid gray line.

M^+^ mutants have some stringent-like transcriptional responses (Rutherford et al., 2009; Zhou and Jin, 1998). Moreover, either lowering the nutritional content of the growth medium or artificially inducing the stringent response leads to high-level resistance to mecillinam and A22 in *E. coli* (Bendezú and de Boer, 2008). We therefore explored whether β-P153L resistance arises from a gene expression program locked into a stringent-like state. We measured differential gene expression in β-P153L and its parental strain without and with induction of the stringent response. We achieved induction of the stringent response by expressing a constitutively active allele of RelA (*relA\**) and compared our results to a published dataset that used the same method (Sanchez-Vazquez et al., 2019) (**Supplemental Dataset 2**).

Steady-state gene expression in β-P153L without *relA\** showed limited similarity to stringent response gene expression. Of the 344 genes significantly differentially expressed in β-P153L and measured in both datasets, only 36% overlapped with the stringent response (**Figure 5C**). Moreover, in the overlapping set, only inhibited genes were significantly enriched (*p*=0.02, odds ratio=1.6 for inhibited genes; *p*=0.09,odds ratio=1.3 for activated genes). Analysis by functional category reinforced the differences between β-P153L expression and the stringent response. By contrast to the uniform repression of ribosomal protein expression during the stringent response, in β-P153L only 8 genes for ribosomal proteins (14%) were significantly repressed and only 5 (8%) were repressed more than 2-fold (**Figure 5D**). Moreover, rather than repressing tRNA-aminoacylation genes, β-P153L significantly activated 12 (46%) (**Figure 5D**). Rather than prevalent activation of amino acid biosynthesis genes, only those for histidine biosynthesis were significantly upregulated in β-P153L (**Figure 5E**). Induction of *relA\** resulted in a response highly overlapping with the classic stringent response in both β-P153L and its parent (**Supplemental Figure 2A, Supplemental Dataset 2**), and the response of the mutant was greater than that of its parent (**Supplemental Figure 2B**), even though the fold induction of *relA\** was similar (**Supplemental Figure 2C**).

Thus, despite *in vitro* behaviors of M^+^ mutants that mimic ppGpp binding (Rutherford et al., 2009; Zhou and Jin, 1998), the steady-state transcriptional program of β-P153L *in vivo* is largely distinct from the canonical stringent response.

### β-P153L protects against death caused by loss of rod shape in rich media

Mecillinam and A22 target PBP2 (Spratt, 1977) and MreB (Gitai et al., 2005), respectively, two components of the cell wall elongation machinery that directs lateral cell wall insertion and maintains rod shape in *E. coli*. They are essential during rapid growth (e.g. in LB), but dispensable in nutrient-poor environments (e.g. M9) (Bendezú and de Boer, 2008). As the stringent response is not obviously responsible for resistance in β-P153L, we sought to understand the origin of resistance by determining the full range of resistance responses and morphological changes associated with growth in the antibiotics.

We compared liquid growth curves in LB for β-P153L and its parental control over a range of A22 and mecillinam concentrations. β-P153L exhibited >10X higher MICs than the control (**Figure 6A**). To investigate how β-P153L protects *E. coli* under A22 or mecillinam treatment, we followed single-cell growth and morphology after exposure to above-MIC concentrations of mecillinam using time-lapse microscopy. Wild-type cells stopped dividing and grew increasingly large, their membranes invaginated, and the cells eventually lysed (**Figure 6B**). By contrast, β-P153L cells morphed from small rods to small cocci that continued to grow and divide (**Figure 6B**). β-P153L cells displayed a similar morphological transition to small cocci in A22 (**Supplemental Figure 3A**), and fluorescent D-amino acid labeling (Kuru et al., 2012; Kuru et al., 2015) during growth with mecillinam revealed that β-P153L cocci retained a cell wall (**Supplemental Figure 3B**), rather than forming cell-wall-less spheroplasts.

**Figure 6:**
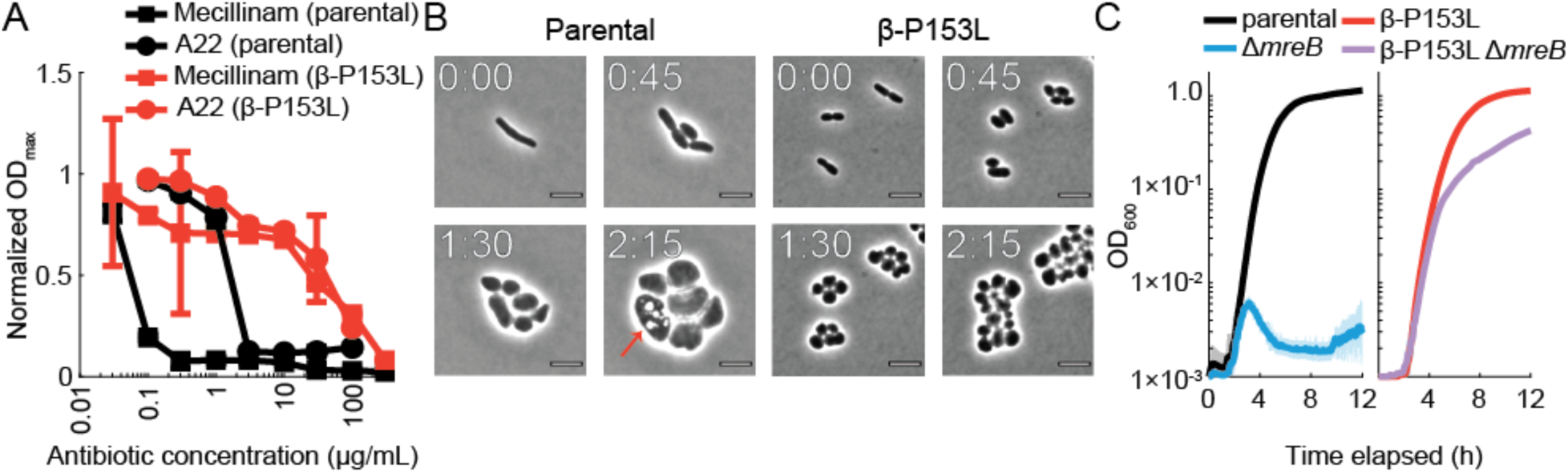
β-P153L renders loss of rod shape non-lethal in rich media. A) β-P153L is highly resistant to both mecillinam and A22, with MICs for both antibiotics increased by at least an order of magnitude. The maximum OD_600_ (OD_max_) was extracted from growth curves of β-P153L (red curves) and its parental control (black curves) and normalized by the ODmax of each strain in the absence of antibiotic. Error bars represent 95% confidence intervals. B) In the parental strain growing on agarose pads with LB+10 µg/mL mecillinam, division rapidly halted and cells expanded dramatically in volume, with the formation of apparent membrane invaginations (red arrows) and eventual lysis. β-P153L cells growing on agarose pads with LB+10 µg/mL mecillinam lost rod-like shape but remained small and continued dividing without lysis. C) β-P153L suppresses the lethality of Δ*mreB* in rich media. The Δ*mreB* deletion was introduced into the background of β-P153L and its parental control under permissive conditions (M9 minimal medium at 30°C). Growth curves were measured after transitioning these strains to non-permissive conditions (LB at 37 °C). Left: Δ*mreB* halted bulk growth after a transition to non-permissive conditions. Right: β-P153L Δ*mreB* retained luxuriant growth in LB.

These results predicted that β-P153L should also render the genes encoding PBP2 and MreB (*mrdA* and *mreB*) non-essential during rapid growth conditions. We constructed Δ*mreB* and Δ*mreB* β-P153L mutants under permissive conditions (minimal medium, 30°C), and tested growth of the double mutant after shifting to non-permissive conditions (LB, 37°C). The Δ*mreB* β-P153L double mutant exhibited essentially normal growth, while the Δ*mreB* control quickly halted growth after the transfer (**Figure 6C**). Whole-genome resequencing confirmed that the strains did not contain second-site suppressors (**Supplemental Table 4**).We conclude that β-P153L renders *mreB* non-essential in rich media by preventing lysis after a loss of rod shape.

### M^+^ mutants have growth rate-independent reductions in cell size

It has been proposed that the irreversible step toward death in A22-treated cells is the expansion of cell width beyond a limit at which division can no longer occur, leading to run-away cell width and eventual lysis (Bendezú and de Boer, 2008). According to this model, the small size of β-P153L cells during treatment could keep the mutant below the non-reversible threshold and prevent death. However, the basis for the small size of β-P153L cells was not immediately clear. *E. coli* and many other rod-shaped bacteria have a log-linear relationship between cell size and growth rate when the nutrient content of the medium is varied (Schaechter et al., 1958; Taheri-Araghi et al., 2015). This relationship, termed the Growth Law, suggested that the smaller size of β-P153L in LB might have been simply due to its lower growth rate.

To test this idea, we measured cell size and growth rate across four media with different nutritional contents. If the small size of β-P153L had been due to a growth rate defect alone, the overall relationship between growth rate and cell size would have been indistinguishable between the two strains. Instead, we found that β-P153L was significantly smaller than its parental strain across all growth rates (**Figure 7A**). To ask whether our conclusions could be generalized to other M^+^ mutants, we chose 6 additional M^+^ mutants and measured the relationship between cell size and growth rate. We found that only a subset of M^+^ mutants had a slow growth phenotype in LB, but all M^+^ mutants had reduced size, with even the most subtle M^+^ mutant exhibiting a 27% reduction in cross-sectional area (**Figure 7B**). We conclude that M^+^ mutants exhibit a spectrum of reduced sizes and that their size reduction is not due solely to slower growth.

**Figure 7:**
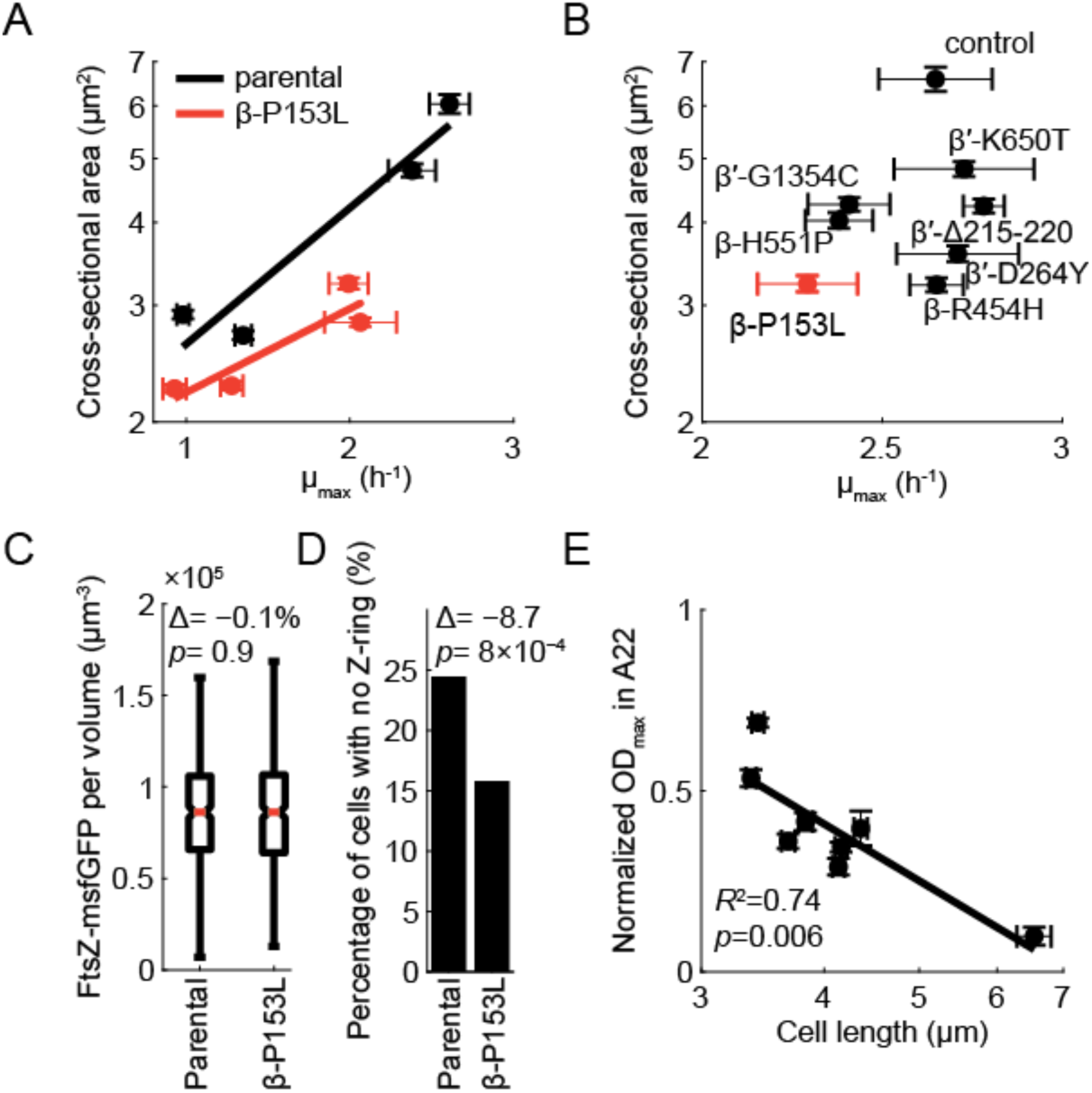
M^+^ mutants stabilize FtsZ to confer A22 resistance. A) β-P153L cells are smaller than the parental control, even after controlling for growth rate. β-P153L and its parental control were for grown multiple generations in log phase in four media: MOPS minimal medium+0.2% glucose, MOPS minimal+0.2% glucose supplemented with 6 amino acids, MOPS complete medium+0.2% glucose, and Tryptic Soy Broth. The maximum growth rate (µ_max_) was extracted from growth curves started with the log-phase cultures. Phase-contrast images of log-phase cells grown in steady were acquired after spotting the cultures on PBS + 1% (w/v) agarose pads, and cell area was computed from the segmented single-cell contours. Straight lines are linear regressions. Error bars on both axes are 95% confidence intervals for individual measurements. B) M^+^ mutant cells are smaller than the parental control. Seven M^+^ mutants from different clusters were grown into log-phase in LB along with their parental control. Cultures were simultaneously spotted onto PBS+1% agarose pads to measure cell size and used to inoculate growth curves to measure maximum growth rate (µ_max_). All M^+^ mutants were significantly smaller than their parental control, while only β-P153L, β-H551P, and β′-G1354C exhibited a statistically significant decrease in µ_max_. C) Total FtsZ-msfGFP fluorescence within each cell contour normalized by the cell volume was not significantly different in β-P153L. The total fluorescence in β-P153L was only 0.1% less than the control (*p*=0.9). Red lines represent population medians, boxes cover the upper and lower quantiles, and whisker lengths are 1.5 times the inter-quartile range. n=500 for β-P153L and n=364 for its parental control. D) The fraction of cells with no detectable FtsZ-ring was significantly lower for β-P153L compared with the parental control (*p*=8×10^−4^, n=500 for cells for β-P153L and n=364 for its parental control). E) A22 resistance is correlated with cell length. The maximum OD_600_ of the 7 M^+^ mutants and their parental control in LB with 13.5 µg/mL A22 was extracted from growth curves and normalized by growth curves of the same strain in LB without antibiotic. Normalized OD_max_ values were strongly correlated with cell length from measurements in (B) (*R*^2^=0.74, *p*=0.006).

### Z-ring stabilization is sufficient for A22 resistance

We noticed that the parental strain of β-P153L, BW25113 *rpoBC-cat*, which differs from the precursor BW25113 by the presence of a chloramphenicol marker between *rpoB* and *rpoC*, exhibited a cell-elongation phenotype during log phase in rich media with longer than normal cells that sometimes filamented to >10 µm. We hypothesize that the elongation phenotype is related to the *cat* insertion as the strain has no other mutations (**Supplemental Table 4**). Given this observation, we reasoned that the smaller size of β-P153L and the other M^+^ mutants could be related to suppression of filamentation.

Interestingly, filamentation in another *rpoB* mutant was previously found to be suppressed by the stringent response (Vinella and D’ari 1994). Moreover, filamentation due to the *ftsZ^ts^* allele *ftsZ84* is suppressed by both induction of the stringent response and an M^+^ mutation in *rpoD* (Powell, 1998), with increased levels of FtsZ84 in both cases (Powell, 1998). Moreover, *mreB* essentiality is suppressed by overexpression of the central components of the division machinery FtsQAZ (Bendezú and de Boer, 2008). We therefore considered whether increased expression of FtsZ underlays both suppression of filamentation and resistance to A22 and mecillinam in β-P153L.

Using single-cell fluorescence microscopy, we quantified the concentration and spatial distribution of an FtsZ-msfGFP translational fusion in β-P153L and its parent. FtsZ concentration was virtually identical (0.1% difference) between the two strains (*p*=0.9, two sample Kolmogorov-Smirnov test) (**Figure 7C**), providing strong evidence that β-P153L does not alter FtsZ concentration. In wild-type cells, the fraction of cells with a detectable FtsZ-ring is directly proportional to nutrient-determined growth rate (Weart and Levin, 2003). Surprisingly, β-P153L increased the population of cells with an FtsZ-ring (**Figure 7D**), despite having a lower growth rate (**Figure 7A**) and the same FtsZ concentration as its parent (**Figure 7C**).

While the filamentous phenotype of the parental *rpoBC-cat* strain indicates that it has the opposite effect of β-P153L, the effects of the β-P153L mutation are clearly distinct from simply correcting for the division deficiency of the parent. The wild-type BW25113 and the parental strain have equivalent MICs for mecillinam and A22 (**Supplemental Figure 4A,B**). Thus, if β-P153L simply corrected the cell elongation phenotype of its parental strain, it would be sensitive to the antibiotics as well. Instead, we propose that β-P153L stabilizes FtsZ-rings independent of FtsZ expression, and that this phenotype suppresses filamentation during normal growth and allows for continued division under the action of A22 and mecillinam.

Since the other 6 M^+^ mutants also exhibited a range of lengths shorter than the parental strain, we sought to test whether cell length was a proxy for FtsZ-ring stabilization that gives rise to A22 resistance. We grew all seven M^+^ mutants in LB with above-MIC concentrations of A22 and mecillinam and compared the normalized maximum OD_600_ in the presence of the drugs to cell size and growth rate parameters. The normalized OD_max_ in the presence of A22 was strongly negatively correlated with cell length (*R*^2^=0.74, *p*=0.006), but not with cell width (*R*^2^=0.14,*p*=0.36) or growth rate (*R*^2^=0.17, *p*=0.31) (**Figure 7E, Supplemental Figure 4C,D).**

Interestingly, resistance to mecillinam had a different origin. The normalized OD_max_ in the presence of mecillinam was correlated with growth rate (*R*^2^=0.53, *p*=0.04) but not cell size (*R*^2^=0.16, *p*=0.32). We conclude that FtsZ-ring stabilization is not sufficient to confer resistance to mecillinam and that another aspect of regulation by the stringent response, likely a correlate of the growth rate phenotype, is necessary. These results are consistent with previous findings that mild FtsZ overexpression is sufficient for survival in A22 but not mecillinam (Cho et al., 2014). We conclude that FtsZ-ring stabilization rather than FtsZ upregulation determines cell length and A22 sensitivity in M^+^ mutants, and that a reduction in growth rate is neither sufficient nor necessary.

## Discussion

As the enzyme responsible for bacterial transcription and the integrator of transcriptional control, RNAP has been the focus of an enormous amount of research. In addition to structural, biochemical, and evolutionary analyses, multiple studies have utilized RNAP-centric genetic approaches, including early work on resistance to RNAP-targeting drugs such as rifampicin (Jin and Gross, 1988) and streptolydigin (Heisler et al., 1993), temperature-sensitive (Saito et al., 1986) and dominant-negative mutations (Sagitov et al., 1993), mutations altering features of RNAP (e.g. attenuation (Weilbaecher et al., 1994), survival in minimal media (Rutherford et al., 2009)), and structure-guided mutational analysis (Wang et al., 2006). In addition to revealing the inner workings of RNAP, this top-down body of work has resulted in novel physiological discoveries. Furthermore, the repeated isolation of RNAP mutations during adaptation to biotechnology-related environments (Cheng et al., 2014; Haft et al., 2014; Tenaillon et al., 2012) has made evident the value and need for a deeper understanding of transcription-related pleiotropy (Alper and Stephanopoulos, 2007).

Here, we show that a bottom-up approach based on unbiased, expansive screening and clustering of the phenotypes of large numbers of RNAP mutations is a powerful tool for functional discovery, illuminating structure-function relationships of RNAP at the single-residue level and systems-level connections between transcription and other cellular processes. That our strategy was successful even though our library is overrepresented in Rif^R^ and M^+^ mutants underscores the fact that mutations isolated under the same selective pressure can have distinct, pleiotropic phenotypes. Importantly, our findings also highlight the patterns behind these pleiotropic fitness trade-offs, hinting at the possibility of general rules that govern the physiological effects of RNAP mutations. We probed one such chemical interaction to discover an effect of M^+^ mutants on cell-size control by modulating FtsZ-ring stability.

Our analysis of the phenotypes of the lineage-specific insertions in the β-subunit highlight both the strengths and challenges of chemical genetic screens. Although neither the β-ΔSI1 nor the β-ΔSI2 strain exhibited chemical sensitivities that clustered with other RNAP mutations, strong chemical sensitivities of β-ΔSI1 were phenotypically correlated with Δ*dksA*. The proximity of β-SI1 to the known binding site of DksA on RNAP allowed us to predict a role for β-SI1 in the binding and function of DksA that was validated with biochemical and genetic evidence (Parshin et al., 2015). Further work by others has fleshed out this interaction by identifying the required conformational changes (Molodtsov et al., 2018), found that additional secondary channel regulators bind to β-SI1 (Chen et al., 2019; Molodtsov et al., 2018), and identified a role of ppGpp in DksA–RNAP interaction (Molodtsov et al., 2018; Ross et al., 2016). By contrast, β-ΔSI2 displayed minor chemical sensitivities (**Figure 4A,B**) and the lack of phenotypic clustering information for β-ΔSI2 prevented hypothesis generation through traditional chemical-genetic inference. Nonetheless, our ability to successfully predict the attenuation-proficient phenotype of the β-I966S mutant in β-SI2 (**Figure 4C,D**), consistent with previous predictions of a hyper-termination phenotype (González-González et al., 2017), highlights the relatively higher predictive power of correlations between mutants for identifying the function of uncharacterized mutations compared with the interpretation of individual sensitivities, as has been the case for most high-throughput genetic screens to date (Schuldiner et al., 2005). Interestingly, the three-mutants comprising the clique within cluster 16 that includes β-I966S are outliers to the negative correlation between phenotypic profile similarity and pairwise distance on the RNAP structure (**Figure 4C**), highlighting the potential for other allosteric interactions in RNAP that remain to be discovered.

Finally, our analysis of A22 and mecillinam resistance in M^+^ mutants (**Figure 5A,B**) demonstrates the power of this approach to generate new insights integrating cell biology and physiology. With prior knowledge that the stringent response confers resistance to A22 (Bendezú and de Boer, 2008) and mecillinam (Bendezú and de Boer, 2008; Vinella et al., 1992), and that M^+^ RNAP enzymes exhibit certain behaviors associated with the stringent response (Rutherford et al., 2009; Zhou and Jin, 1998), we first explored the possibility that A22 and mecillinam resistance reflected stringent-like transcription by M^+^ mutants. However, the transcriptional program of β-P153L was largely dissimilar to the stringent response (**Figure 5C-E**).

Our single-cell analyses revealed that M^+^ mutants are shorter than the parental strain (**Figure 7A,B**) and that β-P153L cells remain as small cocci during A22 or mecillinam treatment (**Figure 6B**), prompting the hypothesis that M^+^ mutants have higher expression of FtsZ. However, FtsZ concentration was unaffected (**Figure 7C**), consistent with previous findings that neither the direct σ^70^ promoters for *ftsZ* (Navarro et al., 1998) nor the upstream σ^S^ promoters (Powell, 1998) responded to elevated levels of ppGpp *in vivo* and that increased expression of FtsZ is neither necessary nor sufficient for mecillinam resistance (Navarro et al., 1998). Instead, we found that β-P153L decreased the fraction of cells without an FtsZ-ring (**Figure 7D**). Moreover, the other 6 M^+^ mutants that we examined showed varying degrees of reduction in cell length relative to the parental strain (**Figure 7B**) that tightly correlated with survival at high A22 concentration (**Figure 7E**). These observations suggest that all M^+^ mutants increase FtsZ-ring stability, although to different extents, and that this molecular phenotype explains both their decreased cell length and their survival of A22. In *E. coli*, OpgH regulates cell length as a function of growth rate by acting as an FtsZ inhibitor during fast growth (Hill et al., 2013).The mechanism by which M^+^ mutants affect FtsZ stability and how this finding relates to current models for growth rate control of cell size (Hill et al., 2013; Vadia et al., 2017) is currently unknown, but it suggests the possibility that there may be mechanisms in addition to OpgH that control cell FtsZ stability. The chemical-genetic and cytological information that we have generated for the M^+^ mutants should provide a powerful resource to investigate this fascinating connection.

The power of this proof-of-principle experiment highlights the value of a bottom-up chemical-genetic approach to interrogating structure-function relationships *in vivo*. In addition to the multiple RNAP-targeting antibiotics with well-characterized resistance determinants that can be used to create mutant libraries through direct selection, the advent of CRISPR editing will remove the classical limitation to mutations with selectable phenotypes, increasing the breadth and power of the initial mutant library. Moreover, the increased capacity of deep sequencing will enable pooled screens in many different conditions to supplement the arrayed screening we utilized here. In addition to RNAP, chemical genetics is ideal for protein complexes and machineries such as cell-wall synthesis machinery, for which extensive libraries of *mreB* and *mrdA* point mutations have recently been created (Shi et al., 2017). Finally, the simplicity of chemical-genetic approaches facilitates the study of RNAP function in a broader set of bacterial species, which could generate fascinating insights into the evolutionary conservation of structure-function relationships and physiological connections of this essential enzyme complex.

## Supporting information

Supplemental Dataset 2 README

## Acknowledgements

We thank Seth Darst for advice on the design of lineage-specific insertion/deletion mutations. We thank Ann Hochschild for sharing mutants in σ^70^ that were used in the screen. We thank Wilma Ross and Rick Gourse for sharing the pALS13 and pALS14 plasmids. We thank Curtis Ross for his contributions to the data browser. Funding was provided by the Allen Discovery Center at Stanford on Systems Modeling of Infection (to K.C.H); C.A.G. was supported by MIRA grant R35GM118061. This research was supported in part by an award from the Department of Energy Office of Science Graduate Fellowship Program (to A.L.S.). K.C.H. is a Chan Zuckerberg Investigator.

## Data availability and code reproducibility

The raw images and *Iris* data files for the chemical genetic screen along with the two datasets generated in this work have been submitted for publication on Dryad. The raw sequencing reads for whole genome resequencing and RNA sequencing have been deposited at NCBI. Reproducible compute capsules have been published on Code Ocean for the major findings and results of this work.

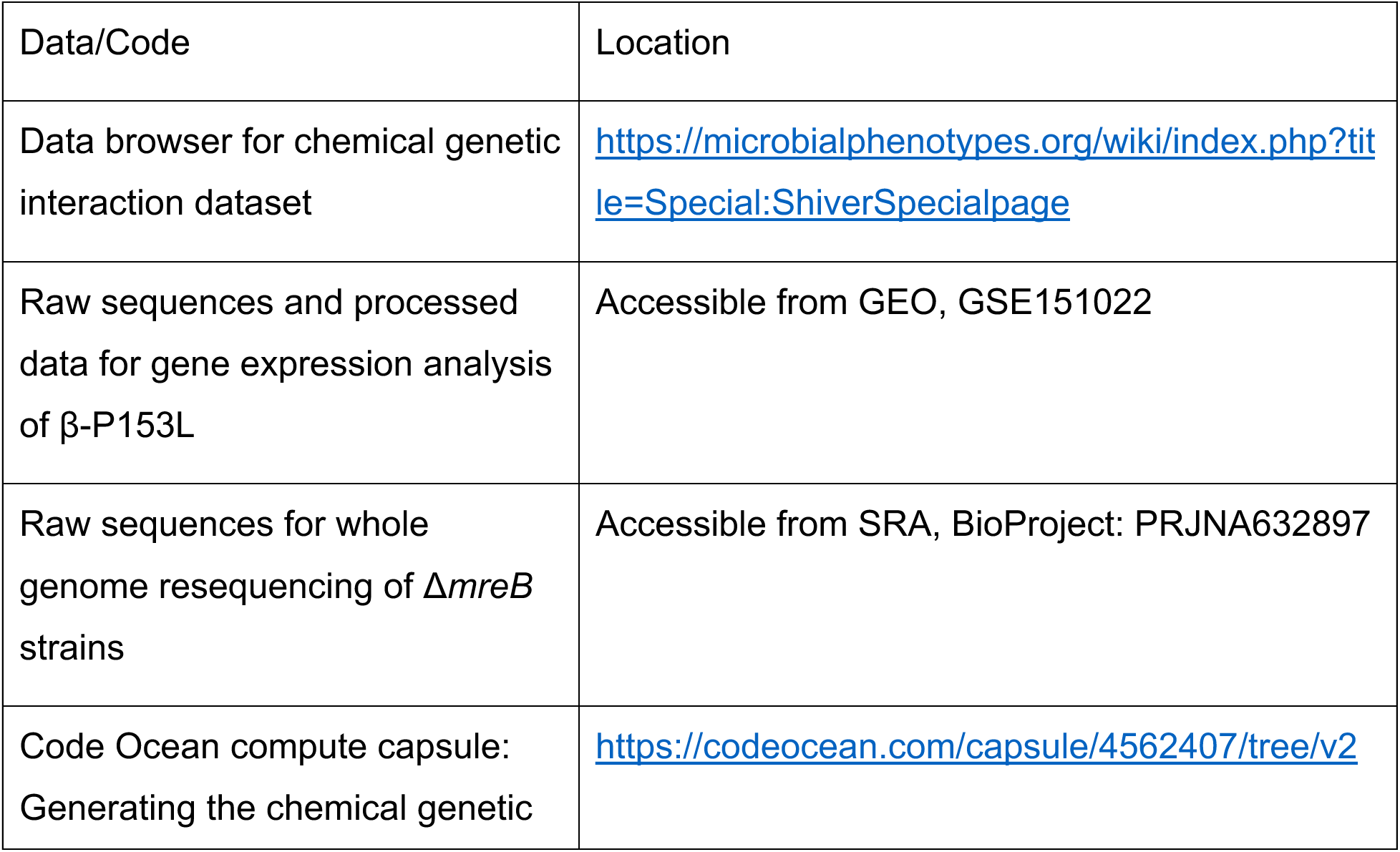
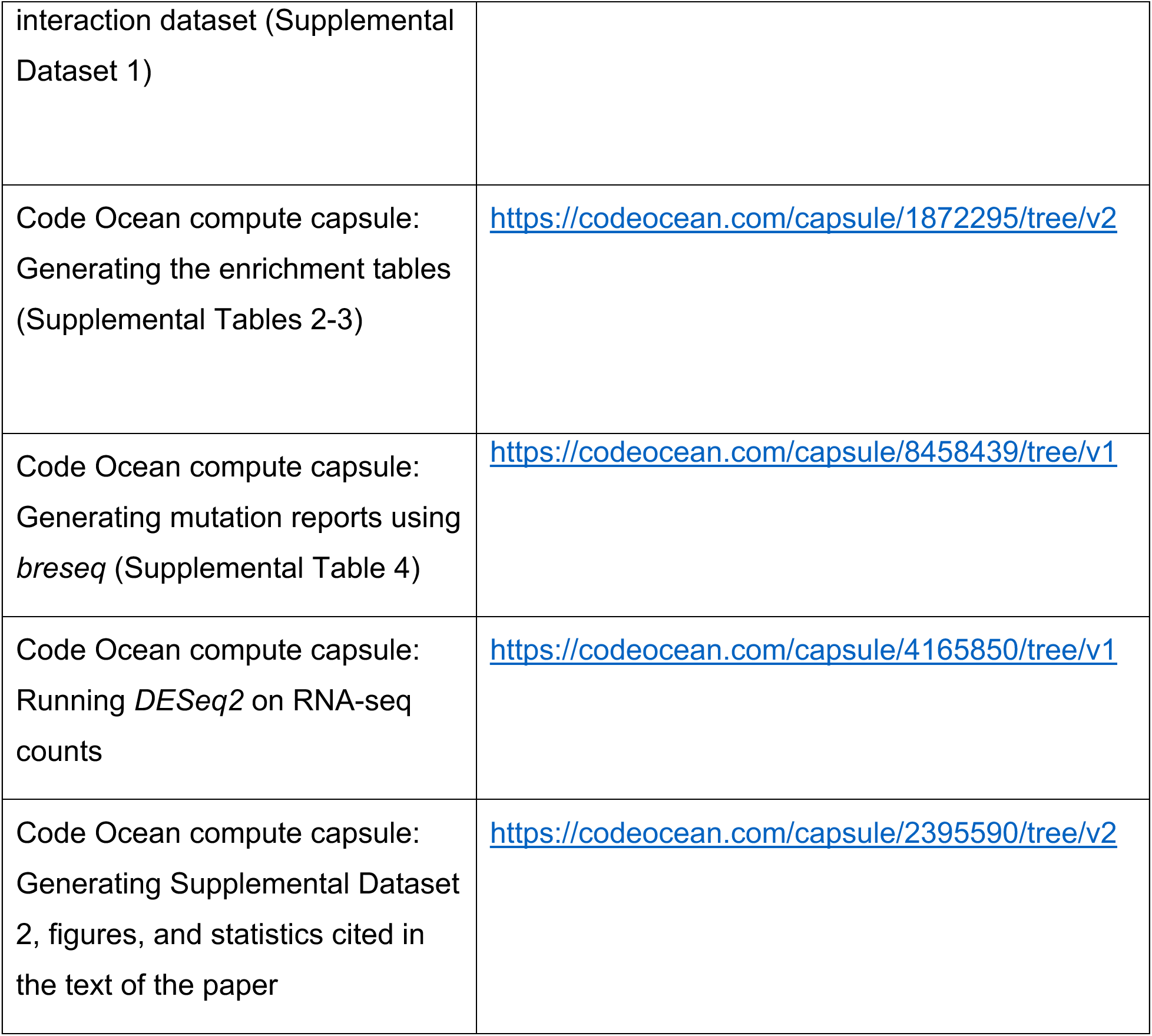

## Methods

### Strains, oligonucleotides, and plasmids

All strains used in this study are listed in **Supplemental Table 1**. All mutations were transduced into or generated in the *E. coli* K-12 BW25113 genetic background for the chemical-genetic screen. During subsequent work to confirm sensitivities found in the screen, we moved some mutations into a MG1655 genetic background using P1*vir* transduction.

All oligonucleotides used in this study are listed in **Supplemental Table 5**. Oligonucleotides were synthesized by Integrated DNA Technologies (Skokie, IL) with standard desalting purification. Oligonucleotides used for recombineering were designed to anneal to the lagging strand to increase efficiency (Ellis et al., 2001). Recombineering oligonucleotides were 79 nucleotides long, unless a high efficiency of mutagenesis was required in which case the length was extended to 89 or 90 nucleotides. For all oligonucleotides, the mismatch(es) were located in the center of the sequence. For oligonucleotides that required highly efficient mutagenesis, four phosphorothioation modifications were included between the five bases closest to the 5’ end of the oligonucleotide to prevent 5’ degradation (Wang et al., 2009).

All plasmids used in this study are listed in **Supplemental Table 5**.

### Mutagenesis using oligonucleotide recombineering

We generated some mutations reported in this study *de novo* using oligonucleotide recombineering. We first transformed strains of the appropriate genetic background with pSIM6 to introduce the λ-Red system (Datta et al., 2006), made electrocompetent cells using published methods (Thomason et al., 2014), and electroporated the cells with mutagenic oligonucleotides. To prepare recombineering-competent cells, an overnight culture was grown at 30 °C in LB with 100 µg/mL ampicillin, and 500 µL of this culture was diluted into 35 mL of fresh LB with 100 µg/mL ampicillin in a 250-mL baffled flask and grown in a shaking water bath (Gyrorotory® Water Bath Model G76, New Brunswick Scientific Co., Incorporated) at 32 °C. The culture was shaken at 330 rpm throughout early log phase until it reached an OD_600_ of 0.4-0.6 as measured on a Genesys 20 spectrophotometer (Thermo-Scientific). Fifteen milliliters of culture were then transferred to a 125-mL baffled flask in an orbital shaking water bath set to 42 °C and 200 rpm for 15 min. After heat shock to induce the λ-Red system from pSIM6, cultures were chilled on ice for 10 min.

Chilled cells were then transferred to a 50-mL conical tube, pelleted at 3,709 xg for 4 min (Allegra X-30R, C0650 adaptor, Beckman Coulter), and resuspended in 50 mL of ice-cold deionized water (MilliQ Biocel A10, Millipore). Cells were pelleted again at 3,709 xg for 4 min and resuspended in 800 μL of ice-cold deionized water. Finally, cells were transferred to a 1.5-mL Eppendorf tube, pelleted in a tabletop centrifuge (Centrifuge 5417 C, Eppendorf) at 10,000 xg for 30 s, and resuspended in 200 µL of ice-cold deionized water. These competent cells were then kept on ice until they were electroporated. Before electroporation, 50 µL of competent cells were mixed with 2 µL of 100 μM oligonucleotide solution before being transferred to an electroporation cuvette. Transformation mixtures were electroporated at 18 kV/cm (Gene Pulser Cuvettes (0.1 cm gap), *E. coli* Pulser, Gene Pulser Attachment, Bio-Rad).

Electroporated cells were immediately resuspended in 500 µL of LB, transferred to a 5-mL test tube, and incubated on a roller drum at 37 °C for 1 h. Recovered cells were then plated according to the selection scheme necessary to isolate the intended mutations.

### Selection of Rif^R^ mutants

Rif^R^ mutations were selected in *rpoB* in BW25113 *rpoBC-cat*. Two hundred microliters of recovered transformants were plated on LB with 10-50 µg/mL rifampicin. Resistant colonies were struck on rifampin plates a second time to purify the colonies and confirm the Rif^R^ phenotype. Mutations were confirmed using Sanger sequencing.

### Selection of M^+^ mutants

We selected for M^+^ mutations in *rpoB* and *rpoC* in BW25113 *rpoBC-cat* Δ*dksA::kan* using a standard genetic selection (Rutherford et al., 2009). After recovery in LB, cells were pelleted at 10,000 xg for 30 s in a tabletop centrifuge and resuspended in 1 mL of M9 minimal medium with 0.2% glucose. Two hundred microliters were then plated on M9 minimal medium plates with 0.2% glucose. Colonies that grew in the first 48 h were struck again on M9 minimal medium plates to confirm growth. Mutations were then transduced into BW25113 using the genetically linked *cat* gene. Separation of the *rpoBC* locus from Δ*dksA::kan* was confirmed by testing for a Kan^s^ phenotype. Co-transduction of the mutations with *cat* was confirmed with Sanger sequencing.

### Screening for attenuation mutants

We screened for attenuation-enhancing mutations in a BW25113 *rpoBC-cat* Δ*trpR::kan* genetic background. Recovered cells were plated in a dilution series on LB agar, and 200 µL of the 10^-3^, 10^-4^, and 10^-5^ dilutions were plated on LB agar plates and grown at 37 °C overnight. Single colonies were patched onto a grid on an LB plate and then replica-plated onto M9 minimal glucose plates supplemented with L-cysteine (400 μg/mL), L-methionine (400 μg/mL), L-leucine (400 μg/mL), indole (5 μg/mL), and 5-methyl anthranilic acid (5-MAA, 100 μg/mL) and grown at 30 °C. Patches with robust growth in the presence of 5-MAA were considered to be potential mutants (Weilbaecher et al., 1994; Yanofsky and Horn, 1981). Single colonies were isolated by streaking from the non-selective patch, and the mutation was confirmed with Sanger sequencing. Finally, the mutant was transduced into BW25113 using the genetically linked *cat* gene. Separation of the mutation from Δ*trpR::kan* was confirmed by testing for a Kan^s^ phenotype. Co-transduction of the mutation with *cat* was confirmed with Sanger sequencing.

### Screening for mutants with PCR

Some of the mutations that we generated using oligonucleotide recombineering had no available selective phenotype, so we directly screened for these mutations using a PCR-based assay. For point mutations, we used the BW25113 *rpoBC-cat* genetic background. We designed oligonucleotides that introduced silent mutations in the codons surrounding the mutation of interest to increase the efficiency of mutagenesis (Thomason et al., 2014) and improve our ability to distinguish between the mutant and wild-type alleles through primer annealing. After transformation with the mutagenic oligonucleotide, recovered cells were plated in a dilution series on LB agar. Single colonies were isolated and suspended in 50 µL of PCR buffer. One microliter of this suspension was used for colony PCR (KAPA2G Robust HotStart PCR Kit, Roche), whereby a ∼500-bp amplicon was amplified by a primer pair in which the 3’ end of one primer was complementary to the mutant allele. A colony that led to amplification with the mutation-specific primers was considered to be a positive hit. To further purify the putative mutant, the colony suspension was struck on LB agar plates, grown overnight at 37 °C, and the PCR screen was repeated. We then verified the mutation of interest using Sanger sequencing.

To generate deletions of the lineage-specific insertions SI1 and SI2, we used the strain BW25113 *rpoBC-cat* Δ*mutS::kan*. Primers were designed to detect the deletions through a shift in amplicon size. After transformation with the mutagenic oligonucleotide, recovered cells were plated in a dilution series on LB agar. Single colonies were isolated and suspended in 50 µL of PCR buffer. One microliter of this suspension was used for colony PCR (KAPA2G Robust HotStart PCR Kit, Roche). Samples with a small amplicon size were considered positive hits. The colony suspension was struck on LB, grown overnight at 37 °C, and the PCR screen was repeated on single colonies. After confirmation of the deletion with PCR, the suspensions were used to inoculate 5 mL of LB, grown to an OD_600_ of ∼0.6, and this culture was used to create a P1*vir* lysate for transduction into BW25113. Co-transduction of the deletions with the *cat* antibiotic resistance gene was confirmed with PCR and then Sanger sequencing. Separation from the Δ*mutS::kan* locus was confirmed by testing for a Kan^s^ phenotype.

### Transduction of existing mutations into the BW25113 background

Some of the mutants used in our study were collected from the scientific community and transduced into a BW25113 background. We first introduced the relevant antibiotic markers into the original strain using λ-Red recombineering with the pSIM6 plasmid. We then transduced the mutations into BW25113 using P1*vir* and selected for the antibiotic resistance cassette that we introduced in the previous step. Co-transduction of the genetically linked mutations was confirmed with Sanger sequencing.

### Design and assembly of the 1536-colony array

We split biological replicates for each transcription mutation or gene deletion into two sets (Array #1 and Array #2). We then arrayed the mutations within each set in triplicate with randomized positions in a 32×48 array of 1536 colonies. To minimize edge effects (French et al., 2016), we filled the outermost two columns and rows of the 1536-colony array with wild-type controls and only analyzed the inner positions. The mutants were split according to antibiotic resistance phenotype (Cam^r^ and Kan^r^) into 16 groups that corresponded to each of the 16 96-well plates that would comprise the 1536 array. Based on the final position in the 1536-well array, spaces in each 96-well plate were devoted to wild-type (either BW25113 or BW25113 *rpoBC-cat*) used as “dummy” colonies that would grow in all conditions.

For storage, plates were grown overnight at 37 °C with shaking at 900 rpm in a humidified platform shaker (Infors HT). Glycerol was added to a final concentration of 12.5%, and aliquots of each plate were stored at −80 °C in a 96-well format. The 2 1536-colony arrays were assembled by thawing copies of the 2 × 16 96-well plates and using a Rotor pinning robot (Singer Instruments) to spot the plates, first into 2 × 4 384-colony plates, and finally into 2 1536-colony format plates (Array #1 and Array #2).

### Screening the library for chemical sensitivities

We pinned Array #1 and Array #2 in parallel onto agar plates with antibiotics added to the agar. The screens were performed in 4 batches with 30 or more conditions in each batch to allow for normalization within each batch (Collins et al., 2010). Chemical perturbations were chosen to overlap with existing chemical genomics datasets (Nichols et al., 2011; Shiver et al., 2016). For the dataset from (Nichols et al., 2011), which used three concentrations per chemical, the concentration with the highest number of significant interactions was chosen. For the dataset from (Shiver et al., 2016), Arrays #1 and #2 were screened at the same time using the same batch of chemicals as used for the gene-deletion library.

For each condition, drug was added to melted LB agar and 45 mL was poured into PlusPlates (Singer Instruments). Source plates were generated by pinning the colony arrays onto LB agar plates, and each source plate was used to pin the array onto multiple drug plates. Plates were incubated at 37 °C for a time interval over which colonies had grown appreciably but had not overgrown to the point that colony edges overlapped. Images were taken with a Powershot G10 camera (Canon) and a custom illumination configuration. Colony opacity was estimated using the software *Iris* v. 0.9.4 (Kritikos et al., 2017).

### Statistical analyses of colony sizes to generate S-scores

Data were analyzed using similar algorithms to the gene deletion library (Shiver et al., 2016), with several specialized steps added to or modifying the analysis. Data in the outer rows and columns were immediately discarded. After normalizing the average colony size on each plate and the spatial bias in colony size due to the amount of colony transferred to the uneven agar surface during the pinning step (Collins et al., 2006), we performed an additional normalization to account for effects in the transcription mutants that were due to the genetically linked antibiotic marker alone. For every control strain (antibiotic marker alone), we computed the multiplicative factor required to make its average colony size equal to the average of the entire plate. We then multiplied the control strain and every associated mutant by this factor to normalize for marker-specific effects.

S-scores were computed for every colony position in the library, and S-scores of the same strain were averaged within each of Array #1 and Array #2 separately. S-scores for each mutant were then averaged between Array #1 and Array #2. Measurements of combinations of mutation and condition for which only one of the two plates passed quality control were transformed by a pseudo-averaging mapping (Collins et al., 2006). Finally, we leveraged the Keio deletion mutants included in our screen to compare our results to a genome-wide deletion dataset (Shiver et al., 2016) on a condition-by-condition basis. We used the S-scores to calculate a correlation between each condition in the current work and in the gene deletion dataset. We split the cross-dataset comparisons into matched pairs (same stress condition label, any concentration) and non-matched pairs (all other comparisons). Conditions were retained in the analysis only if the correlation from the matched pairs was greater than the 95^th^ percentile of all non-matched correlations.

### Defining and visualizing significant clusters

Initial attempts to define the optimal number of clusters for k-means clustering of the dataset using the elbow method were unsatisfactory, so we developed an alternative approach.

In the first step, we hierarchically clustered a randomized copy of the dataset. We then calculated the smallest cophenetic distance in the randomized dendrogram. After repeating these steps 30,000 times, we used the 5^th^ percentile of the distribution of cophenetic distances as a cutoff to define significant clusters in the original dataset. The cutoff represents the cophenetic distance that is closer than the closest distance in 95% of randomized matrices. Linkage was calculated using average correlation and distance was calculated using correlation. To speed up iteration over randomized matrices, missing values were imputed with a zero value. The matrix was randomized using *shake* v.5.0 from MATLAB File Exchange (Jos(10584), 2020).

Notably, the distance between cluster 14 and 15 was very close to the cutoff used to define statistically significant clusters. Estimating the cutoff using fewer iterations (∼1,000) led to run-to-run variation in the determination of whether cluster 14 and cluster 15 were separated; increasing the iterations stabilized their separation.

The full dendrogram along with the statistically significant clusters is shown in **Supplementary Figure 1**. The undirected graphs in **Figure 2** were generated by exporting mutant-mutant correlations that exceeded the cutoff into *Cytoscape* v. 3.7.2 (Shannon et al., 2003).

### Generating enrichment tables

Enrichment tables were created by combining two metrics for the significance of chemical genetic interactions. First, we calculated the significance of individual S-scores based on the total distribution of S-scores in each condition (Nichols et al., 2011). Second, we calculated the significantly enriched chemical perturbations in groups of genes defined either by clustering or by previous knowledge. We then combined this information to identify chemical perturbations with S-scores that were enriched for either positive or negative values in a group of mutations that also had at least one individually significant interaction within the group.

We calculated the enrichment of S-scores within mutant groups using the software *Gene Set Enrichment Analysis* v. 3.0 (Subramanian et al., 2005). Ranked lists of S-scores for all transcription mutants were exported for every individual screen condition. Groups were defined from hierarchical clusters and from predefined classes such as ethanol-tolerant mutations or M^+^ mutations. GSEA was run on every individual condition to look for enrichment of the mutant groups in either positive or negative S-scores. Options for GSEA included running the analysis for pre-ranked lists and the flags “-norm meandiv”, “-scoring_scheme weighted”,“-create_svgs false”,“-make_sets true”, “- plot_top_x 20”, “-rnd_seed 081889”,“-set_max 500”,”-set_min 3”,”-zip_report true”, and “-gui false”. The output for each condition was then collected into a single table and the combined set of nominal *p*-values was corrected using the Benjamini-Hochberg FDR correction. If a chemical condition was significantly enriched within a group (adjusted *p* < 0.05) and had at least one individually significant chemical interaction within the group, it was denoted as a significant enrichment. **Supplemental Table 2** contains the enrichment tables for hierarchical clusters. **Supplemental Table 3** contains the enrichment tables for previously defined classes.

### Liquid growth curves

Growth curves were measured in a Synergy H1 (BioTek Instruments) or an Epoch 2 (BioTek Instruments) plate reader using *Gen5* v. 3.04 (BioTek Instruments). Data was collected for approximately 24 h at 37 °C using a 2-min discontinuous loop comprised of a read step at 600 nm and 1 min of a double-orbital shaking step at slow orbital speed and an orbital frequency of 237 cycles per minute.

All experiments used clear, flat-bottom, polystyrene 96-well plates (Greiner Bio-one) covered with a clear polystyrene lid (E&K Scientific). All conditions other than ethanol used a final culture volume of 200 μL per well. The ethanol experiments used a final culture volume of 150 μL per well and 50 μL of mineral oil was overlaid on the culture to reduce evaporation of ethanol.

All cultures to measure chemical sensitivities were inoculated at an OD_600_ of 0.01. The growth curves of Δ*mreB* strains were inoculated at an OD_600_ of 10^-4^ because we found that a lower inoculation density clarified the growth defects of MG1655 Δ*mreB*. For growth in mecillinam, A22, and minimal medium, the inoculum was log-phase culture that had been kept below an OD_600_ of 0.3 using sequential back-dilutions in the same medium as the growth curve for 6-8 h. For transition experiments of Δ*mreB* strains, the inoculum was log-phase culture that had been kept at a low OD_600_ using sequential back-dilutions in M9 minimal medium with 0.2% glucose at 30 °C. For sensitivity to ethanol, hydroxyurea, and trimethoprim, the initial inoculum was a stationary-phase culture that had grown for 16-24 h in LB. Stationary-phase cultures were used in these measurements to enhance sensitivity of β-ΔSI2 to the compounds.

### Quantification of ethanol, trimethoprim, and hydroxyurea sensitivity

The β-ΔSI2 strain did not have an appreciably different minimum inhibitory concentration, maximum OD_600_, or maximum growth rate from the parental control in any of the conditions measured. Instead, growth of β-ΔSI2 slowed at an earlier OD_600_ value at lower concentrations of the compounds. To quantify this effect, we measured area under the curve (AUC). To define a range for the AUC, we used the growth curve of each strain with no drug as a reference point. We extracted the maximum growth rate from the curve, computed the time at which the growth rate first dropped below 10% of this value, and added two hours to define the time *t*_2_ that determines the upper limit of the area to be measured for every drug concentration for that strain. This calculation sets the upper limit for the area of integration to two hours into the transition phase of the strain when grown without stressors. The lower limit of the area to be integrated was the initial time *t*_1_ that measurements started. The area integrated was the blanked OD_600_ between times *t*_1_ and *t*_2_. The OD_600_ was not log-transformed for this calculation. We then normalized the mean AUC for every combination of strain and drug concentration to the no-drug control. The no-drug control was included in every plate and every measurement was compared to the control on the same plate.

### trp-locus attenuation assay

We used the *trp*-locus attenuation assay from (Weilbaecher et al., 1994; Yanofsky and Horn, 1981) to test for a hyper-attenuation phenotype. We transduced mutations and controls from the BW25113 background to an MG1655 Δ*trpR* genetic background using P1*vir* transduction and selected for the linked *rpoBC-cat* antibiotic resistance cassette. The sequences of all transductants were verified with Sanger sequencing.

We used resistance to 5-MAA to test for hyper-attenuation at the *trp* locus. Strains were grown overnight in M9 minimal medium with 0.2% glucose at 30 °C, pelleted using centrifugation, and resuspended at a normalized OD_600_ of 1.0 in M9 minimal salts. We then spotted 2.5 μL of the resuspended cultures onto M9 minimal medium agar plates supplemented with 0.2% glucose, L-cysteine (400 μg/mL), L-methionine (400 μg/mL), L-leucine (400 μg/mL), and indole (5 μg/mL) to which 5-MAA had either been excluded (−) or added at a concentration of 100 μg/mL (+). The spots were allowed to grow at 30 °C for 2 days before pictures were taken with an EOS Rebel T5i (Canon).

### Sample preparation for RNA-seq of β-P153L

The parental *rpoBC-cat* and β-P153L strains were first transformed with the pALS13 (*Ptrc::relA\**) and pALS14 (*Ptrc::relA-*) plasmids. Cells were grown overnight in Teknova Rich Defined Media (EZ-RDM) with 100 µg/mL ampicillin to maintain plasmid selection. Overnight cultures were inoculated into fresh EZ-RDM with 100 µg/mL ampicillin to maintain plasmid selection. After the strains had grown to mid-log phase, samples were taken for the uninduced control, 10 mg/mL isopropyl β-d-1-thiogalactopyranoside (IPTG) was added to induce expression of the *relA* alleles, and samples were taken 5 min after induction. All samples were immediately stored on ice with a 1:8 volume of 5% phenol in ethanol as a stop solution. Samples were transferred to a −80°C freezer for storage before further processing.

### RNA isolation and library prep

RNA was isolated from frozen cell pellets using Trizol (Invitrogen) extraction according to the manufacturer’s protocol. One microgram of purified RNA was fragmented at 95 °C for 7 min in 1X T4 RNA Ligase buffer (NEB) with an equal volume of 2X alkaline fragmentation buffer (0.6 volumes of 100 mM Na_2_CO_3_ plus 4.4 volumes of 100 mM NaHCO_3_). After 3’-end healing with PNK (NEB) in T4 RNA ligase buffer for 1 h, 3’ ligation to a pre-adenylated, barcoded TruSeq R1 adapter with 5 random bases at its 5’ end was performed overnight. The barcoded samples were then pooled and run onto a 6% TBE-Urea gel for size selection (>15 nucleotide insert size), eluted, and ethanol precipitated before performing ribosomal RNA subtraction (RiboZero). Reverse transcription with SuperScript IV (Invitrogen) was performed using a TruSeq R1 RT primer, followed by ligation of the TruSeq R2 adapter to the 3’ end of the cDNA overnight, prior to another gel size selection as described above. A final PCR of the library was performed with indexed TruSeq PCR primers to add the index and P5/P7 flowcell adapters, followed by gel extraction, precipitation, and a BioAnalyzer (Agilent) run for quality control before sequencing on a HiSeq4000 platform.

### Generating and visualizing the differential gene expression datasets

The indexed raw sequencing data was demultiplexed according to their R1 barcodes and the degenerate linker sequence was clipped using a custom script. Mapping of individual reads to the *E. coli* genome (GenBank ID U00096.3) was performed with STAR (Dobin et al., 2013), followed by read counting for individual genome regions according to gene annotations from assembly ASM584v2.

Raw read counts for all 24 samples were used as input for *DESeq2* v. 1.22.2 (Love et al., 2014). Information on the *strain* (parental or mutant), *plasmid* (pALS13 or pALS14), and *induction* (yes or no) was used to group samples for statistical analysis.

Estimates for the differential expression of genes in response to induction of RelA* were obtained using the full factorial linear model *y ∼ strain + plasmid + induction + strain:plasmid + strain:induction + plasmid:induction + strain:plasmid:induction*.

To estimate the response of the parental strain with pALS14 to induction of RelA*, the reference levels for the three factors were set to (*strain*: parental), (*plasmid*: pALS14) and (*induction*: no). The (*induced*: yes) versus (*induced*: no) contrast was used as output.

To estimate the response of the parental strain with pALS13 to induction of RelA*, beyond the response of the parental strain with pALS14, the same model and reference factors were used, but the interaction term of *plasmid*:*induction* was used as output.

To estimate the response of the mutant strain with pALS13 to induction of RelA*, the same model and reference factors were used as above, but the interaction term of *plasmid*:*induction* was used as output.

To estimate the response of the mutant strain with pALS13 to induction of RelA* beyond the response of the parental strain with pALS14, the same model and reference factors were used, but the interaction term of *plasmid*:*induction* was used as output.

Finally, to estimate the effect of the mutant strain alone, we used the average contrast between mutant and wildtype for strains with pALS14 across induced and uninduced conditions. The linear model used was *y* ∼ *strain* + *plasmid* + *induction* + *strain*:*plasmid* + *strain*:*induction*. The reference factors were set to (*strain*: parental), (*plasmid*: pALS14), and (*induction*: no). The (*strain*: mutant) versus (*strain*: parental) contrast was combined with 50% of the *strain*:*induction* interaction term as output.

Before counting the overlap between β-P153L and the *relA\** condition, we filtered both datasets so that they only included genes that were measured in both experiments. The list of significant genes was combined with a flag indicating the direction of change (activated or repressed) and this modified gene set was used as an input for generating the Venn diagram in **Figure 5A**. Intersections between sets were calculated and used as an input to the Venn package in MATLAB File Exchange (Darik, 14 Feb 2011).

### Quantification of A22 and mecillinam sensitivity

Certain combinations of mutants and growth conditions led to a clear shift in the minimum inhibitory concentration of both A22 and mecillinam. To quantify this shift, the maximum OD_600_ (OD_max_) of each culture was computationally extracted from growth curves. To account for plate-to-plate, reader-to-reader, and day-to-day variability in OD_600_ measurements, as well as the idiosyncratic growth curves of the strains and growth conditions, the OD_max_ of every combination of strain and drug concentration was normalized by a no-drug control that was run within the same plate. We calculated the ratio of the means of OD_max_ values at every drug concentration normalized by the no-drug control.

### Fluorescent D-amino acid incorporation

Fluorescent D-amino acid labeling of the cell wall was performed according to the protocols of (Kuru et al., 2015). β-P153L was grown into log phase in LB broth and transferred to LB broth with 30 μg/mL mecillinam. The strain was grown for two doublings in the presence of mecillinam, and back diluted to an OD_600_ of 0.05 in LB broth with 30 μg/mL mecillinam and 500 μM HADA and grown for 1.5 h. It was then washed three times in phosphate buffer saline, and 1 μL was spotted onto phosphate buffer saline 1% (w/v) agarose pads. Fluorescence microscopy images were collected using a Ti-E microscope (Nikon) with a 100X (NA: 1.4) objective and a Zyla 5.5 sCMOS camera (Andor).

### Whole genome resequencing of Δ*mreB* strains

The raw reads from whole genome resequencing were used as input into *breseq* v.0.33.2 (Deatherage and Barrick, 2014) to elucidate potential mutations.

### Generating plots of growth rate versus cell size

For the data in **Figure 7A**, we individually inoculated the parental control (CAG67202) and β-P153L (CAG68095) into 5 mL test tubes filled with one of four media (8 total cultures). The media used were MOPS minimal medium+0.2% glucose, MOPS minimal medium+0.2% glucose supplemented with 6 amino acids, EZ-RDM (Teknova), and Tryptic Soy Broth. For the data in **Figure 7B**, we individually inoculated 8 strains into 5 mL test tubes filled with LB (8 total cultures). The remaining steps were equivalent for both datasets.

After incubating for ∼16 h at 37 °C on a roller drum, each culture was back diluted 1:200 into 3 mL of pre-warmed (37 °C) medium of the same type in 5 mL tubes and incubated in a roller drum at 37 °C. All cultures were continuously monitored and back diluted into pre-warmed media over 6 h to ensure that even the slowest growing cultures had grown in log phase long enough for cell size to stabilize.

After growing all cultures into log phase, each culture was split into two experiments. In the first, culture densities were normalized to an OD_600_ of 0.1, then used to inoculate the same media in a 96-well plate at a final volume of 200 µL and an initial inoculum with an OD_600_ of 0.01. Growth curves were measured as described above, and maximum growth rates were computationally extracted from the growth curves. For the data in **Figure 7E** and **Supplemental Figure 4C,D**, in addition to growing the M^+^ mutants in LB, we also generated liquid growth curves in LB with 13.5 µg/mL of A22 or mecillinam. The maximum OD_600_ was computationally extracted from the growth curves in the presence of drug and normalized against the maximum OD_600_ of the same strain in LB without antibiotic.

In the second experiment, cultures were directly spotted onto a phosphate buffer saline 1% (w/v) agarose pad and phase contrast images were acquired using a Ti-E microscope (Nikon) with a 100X (NA: 1.4) objective and a Zyla 5.5 sCMOS camera (Andor). Phase-contrast images were segmented and meshed using *Morphometrics* (Ursell et al., 2017) and shape parameters were computationally extracted from the mesh.

### Single-cell quantification of FtsZ abundance and localization

P1 transduction was used to introduce the *ftsZ-msfgfp::kan* allele from KC358 into the parental *rpoBC-cat* and β-P153L strains. To quantify FtsZ-msfGFP, overnight cultures grown in LB were inoculated into fresh LB by adding 50 µL of culture to 5 mL of fresh medium. Strains were kept in log phase via serial back-dilution for 5 h. After growth in log phase, cells were harvested at an OD_600_ of 0.06-0.12. Cultures were plunged into ice and 500 µL of 1 mg/mL erythromycin was added to stop further expression of FtsZ-GFP. One microliter of this mixture was spotted on a phosphate buffer saline 1% (w/v) agarose pad and images were collected. Phase and fluorescence microscopy images were collected using a Ti-E microscope (Nikon) with a 100X (NA: 1.4) objective and a Zyla 5.5 sCMOS camera (Andor).

Phase-contrast images of cells were segmented using *DeepCell* (Van Valen et al., 2016) and the cell periphery feature was used to generate cell contours using *Morphometrics* (Ursell et al., 2017). We background-corrected the fluorescence images by subtracting the median value of pixels that were not contained in a contour as defined above from the entire image and calculated the mean fluorescence as the sum of background-subtracted fluorescence within each contour divided by the calculated volume. Cell volume was estimated as series of cylinders with parameters defined by the pill mesh. We filtered the dataset to eliminate poorly fit contours.

To detect FtsZ-rings, we quantified fluorescence along the inner face of cell contours using in-house Matlab scripts and detected peaks with a minimum prominence. Cell contours with less than two peaks were counted as lacking an FtsZ-ring.

## Figure Legends

**Supplemental Figure 1:**
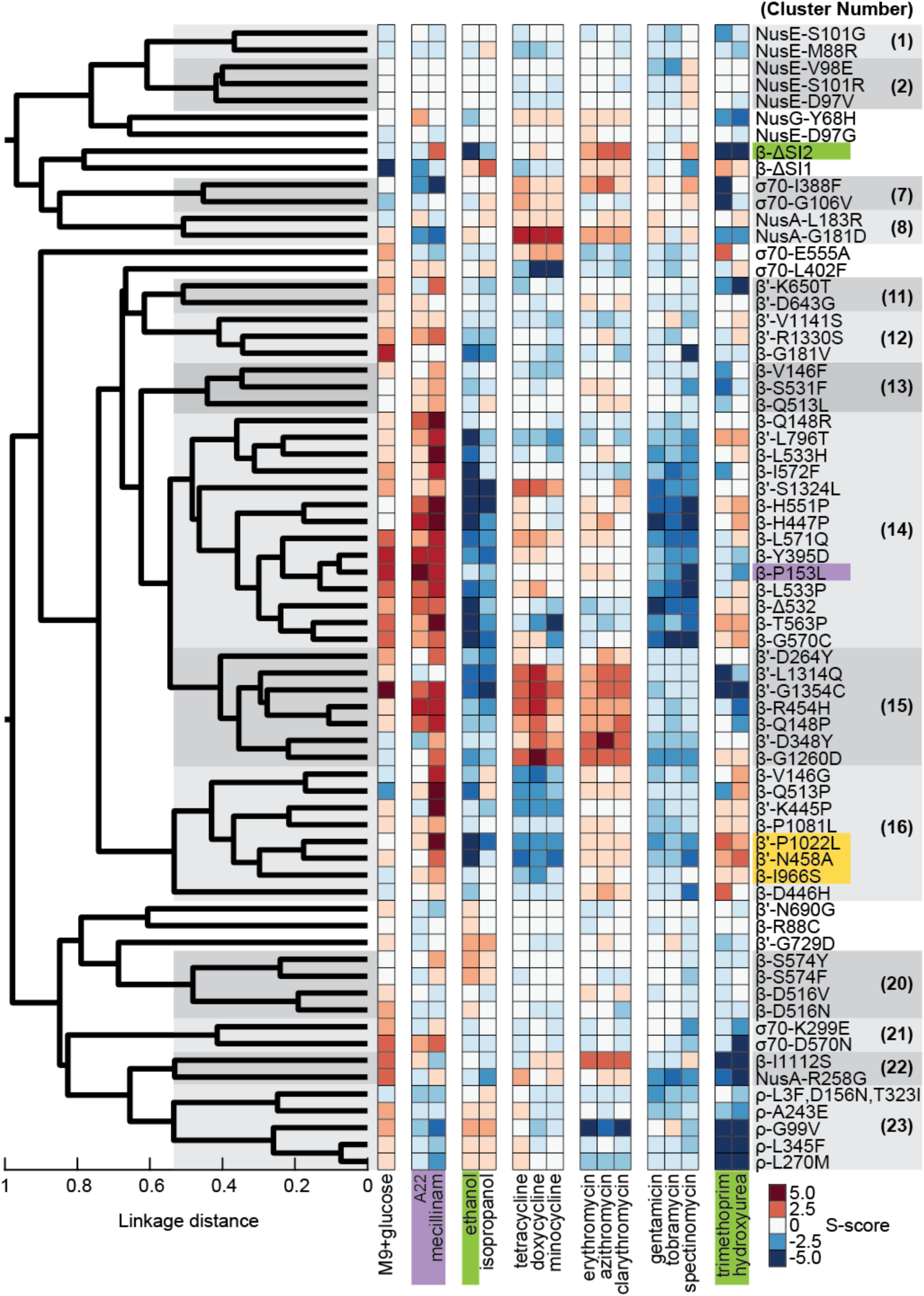
Dendrogram of significant hierarchical clusters and a heatmap of relevant chemical-genetic interactions. Transcription mutants were hierarchically clustered according to shared sensitivities in our chemical-genetic dataset. A cutoff for linkage distance (Methods) identified 23 clusters within the dataset. A subset of the chemical conditions that had significant enrichments among the transcription mutant clusters were chosen for further analysis. A22 and mecillinam (second column group from the left, purple) had widespread positive S-scores that were concentrated in, but not limited to, cluster 14 and cluster 15. Alcohol treatments (third column group from the left) had widespread negative S-scores that were concentrated in, but not limited to, cluster 14 and cluster 15. Sensitivity to select aminoglycosides (gentamicin, tobramycin, spectinomycin, sixth column group from the left) was enriched in cluster 14 (Figure 3B). Resistance to tetracycline (tetracycline, doxycycline, and minocycline, fourth column group from the left) (Figure 3C) and macrolide (erythromycin, azithromycin, clarithromycin, fifth column group from the left) antibiotics was enriched in cluster 15. Sensitivity to tetracyclines was enriched in cluster 16 (Figure 3C). Sensitivity to trimethoprim and hydroxyurea (right-most column group) was enriched in cluster 22 and cluster 23. Resistance to trimethoprim and hydroxyurea was enriched in cluster 16 (Figure 3D). Sensitivity of β-ΔSI2 to ethanol, trimethoprim, and hydroxyurea (green) was confirmed using liquid growth curves (Figure 4A,B). The high correlations among β′-P1022L, β′-N458A, and β-I966S (yellow) were used to predict a hyper-attenuation phenotype for β′-N458A and β-I966S (Figure 4C,D). Resistance of β-P153L to mecillinam and A22 (Figure 5A) was confirmed using liquid growth curves (Figure 5B).

**Supplemental Figure 2:**
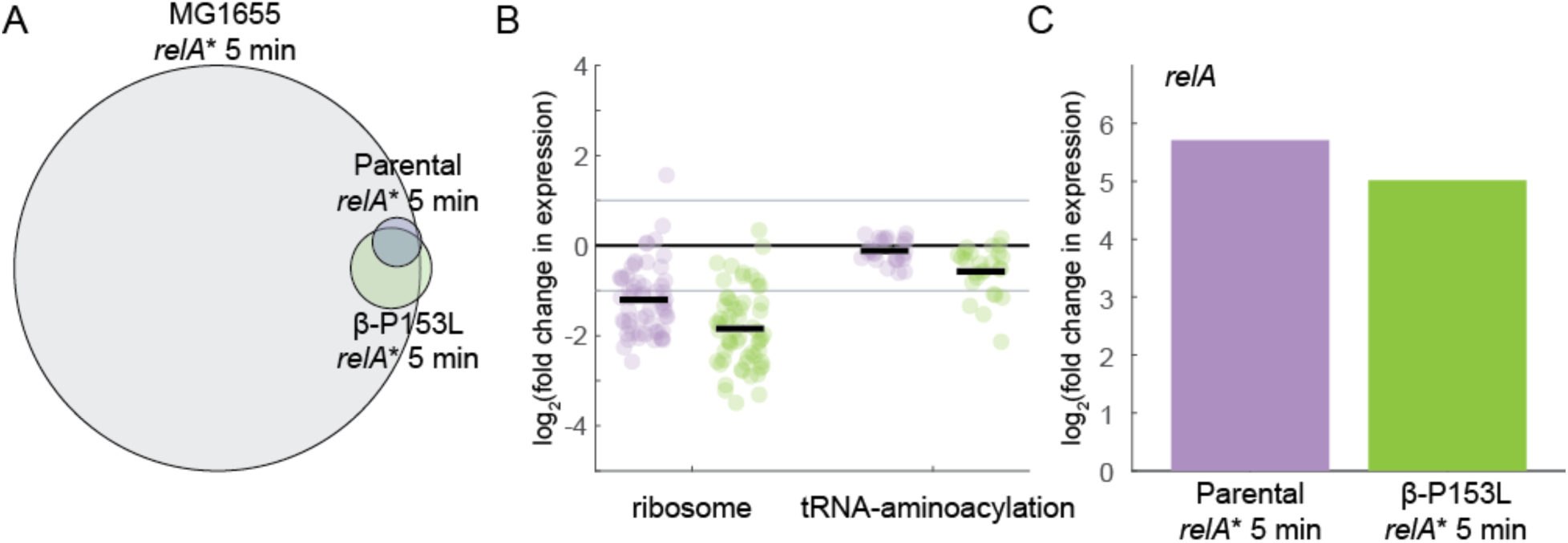
Induction of *relA*\* in β-P153L leads to repression of translation-related genes. A) There is extensive overlap in the significantly differentially expressed genes due to induction of *relA*\* for 5 min in β-P153L (green, data collected in this study), the parental strain (purple, data collected in this study), and MG1655 (gray, from (Sanchez-Vazquez et al., 2019)). The reference dataset (gray) has a more extensive response than the experiments in this study. B) β-P153L (green) and its parental strain (purple) experienced repression of ribosome expression-related (ribosome) and tRNA aminoacylation-related genes after induction of *relA*\*. These data are consistent with the effects of the stringent response. The fold-changes were more pronounced in β-P153L than in the parental strain. Individual genes are plotted as circles. The set average is shown as a sold black line. 2-fold changes in expression are marked with a solid gray line. C) The stronger effects of *relA*\* induction in β-P153L (B) were not due to higher expression of *relA*\*. Despite having a stronger repressive effect on translation-related genes, *relA* induction was slightly lower in β-P153L compared to its parental strain.

**Supplemental Figure 3:**
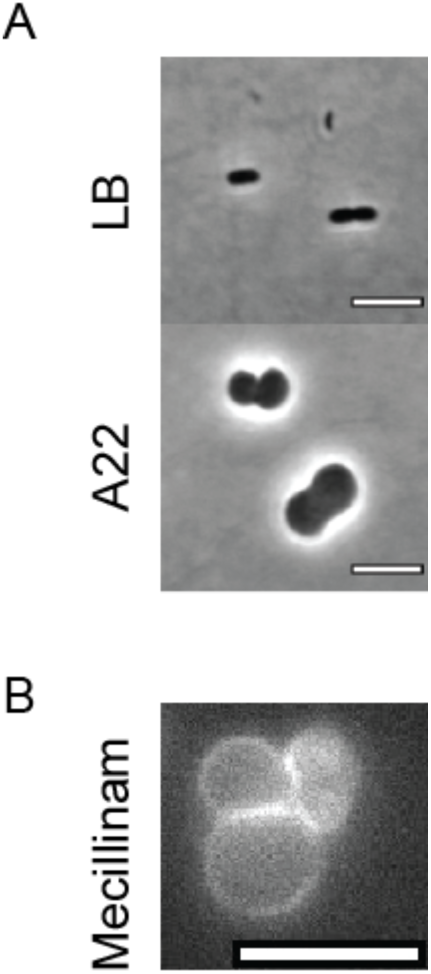
β-P153L grows as cocci in A22 and still has a cell wall during mecillinam treatment. A) Although β-P153L prevents death due to A22 treatment, β-P153L cells still lose rod shape in the presence of A22. Representative phase-contrast images of log-phase β-P153L cells in LB (top) and LB+15 µg/mL A22 (bottom). Scale bar: 5 µm. B) β-P153L cells treated with mecillinam still have a cell wall. Log-phase β-P153L cells grown in the presence of 15 µg/mL mecillinam were pulse-labeled with the fluorescent D-amino acid analogue HADA. Incorporation of HADA into the cell periphery demonstrated that coccoidal β-P153L retains its cell wall during mecillinam treatment. Scale bar: 5 µm.

**Supplemental Figure 4:**
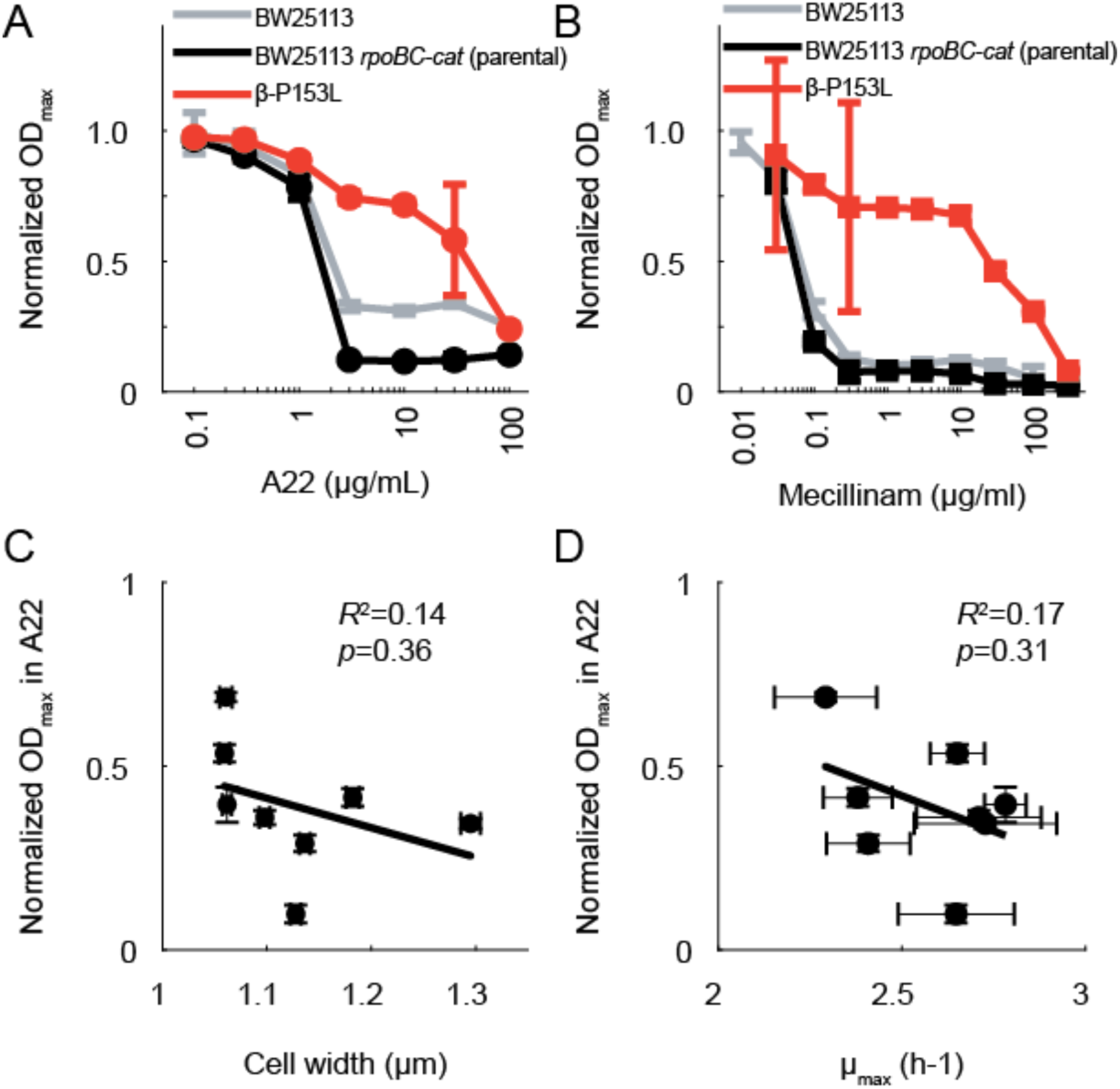
Resistance of β-P153L to mecillinam and A22 shows that it does not simply correct the cell elongation phenotype of its parental strain, and A22 resistance among M^+^ mutants is not correlated with either cell width or growth rate. A) β-P153L is highly resistant to A22 while both its parental strain (BW25113 *rpoBC-cat*) and BW25113 are susceptible. The maximum OD_600_ (OD_max_) was extracted from growth curves of β-P153L (red), its parental control (black), and BW25113 (gray) and normalized by the OD_max_ of each strain in the absence of antibiotic. The growth medium for all 3 strains was LB. Error bars represent 95% confidence intervals. Data for β-P153L and the parental strain are identical to those in Figure 6A and are replotted here for comparison to BW25113. The parental strain halts growth earlier than BW25113 when challenged with A22, but the concentration at which A22 becomes lethal is identical between the two within the resolution of this experiment. By contrast to both wild-type strains, β-P153L was highly resistant to A22. B) β-P153L is highly resistant to mecillinam while both its parental strain (BW25113 *rpoBC-cat*) and BW25113 are susceptible. Data is as in (A), except strains were challenged with mecillinam instead of A22. The concentration at which mecillinam became lethal was identical between BW25113 and the parental strain within the resolution of this experiment. By contrast to both wild-type strains, β-P153L was highly resistant to mecillinam. C) Resistance to A22 was not correlated with cell width. The maximum OD_600_ of the 7 M^+^ mutants and their parental control in LB+13.5 µg/mL A22 was extracted from growth curves and normalized by growth curves of the same strain in LB without antibiotic. The normalized OD_max_ of the 7 mutants was not correlated with the average cell width of log-phase cells grown in LB (*R*^2^=0.14, *p*=0.36). D) The normalized OD_max_, calculated as in (C) was not correlated with the maximum growth rate (µ_max_) extracted from growth rates of the mutants in LB without A22 (*R*^2^=0.17, *p*=0.31).

## Tables

**Supplemental Table 1:**
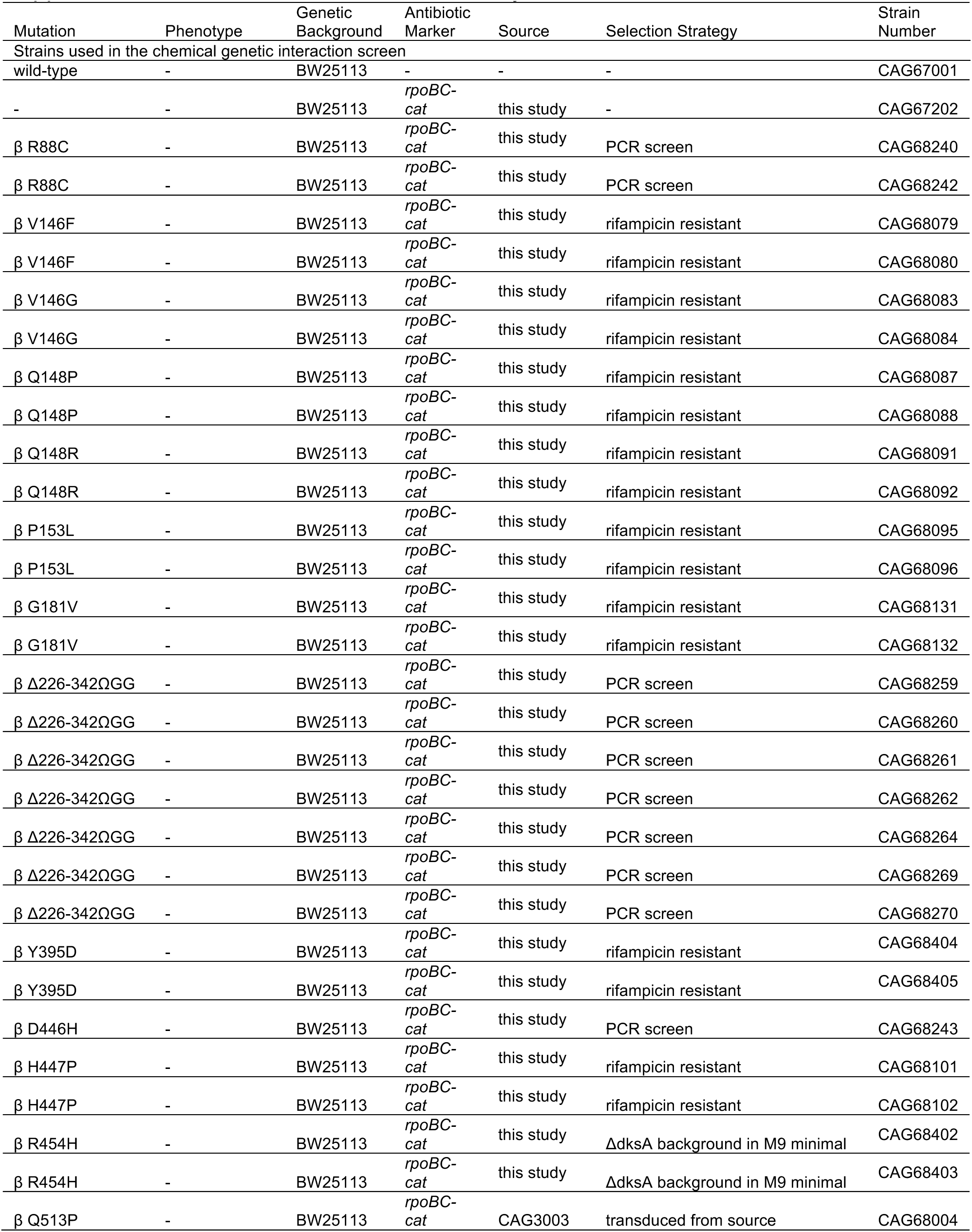
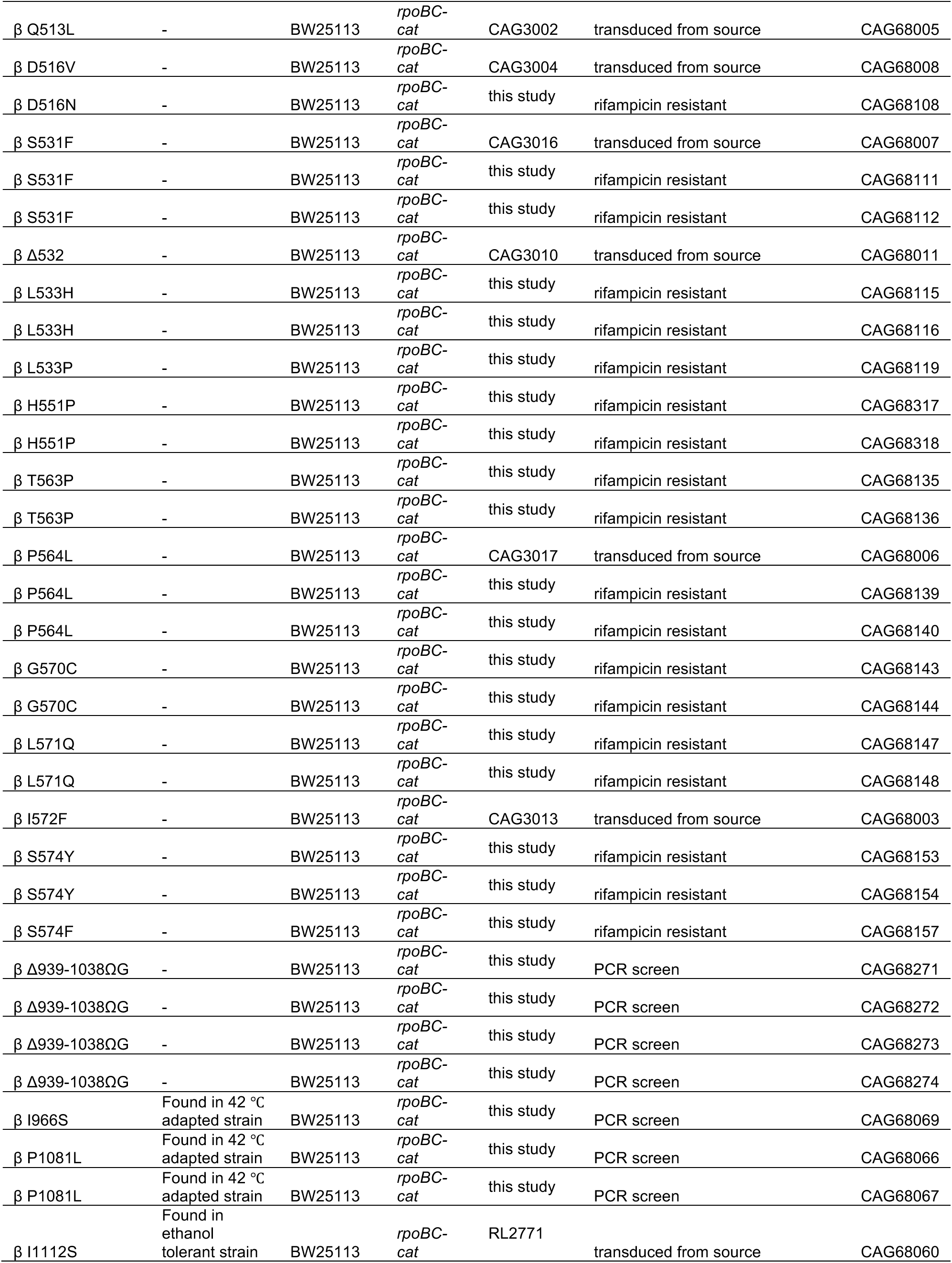
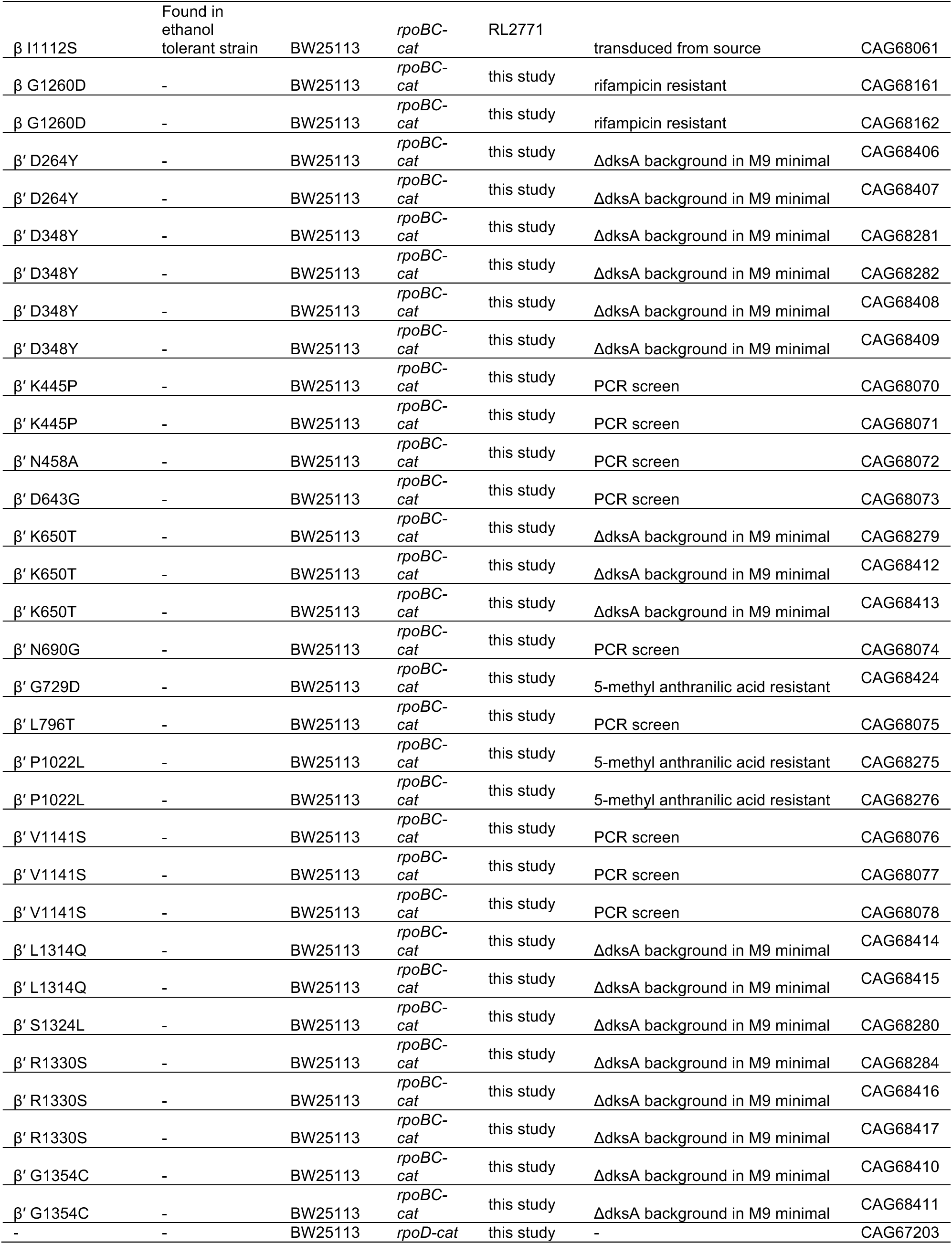
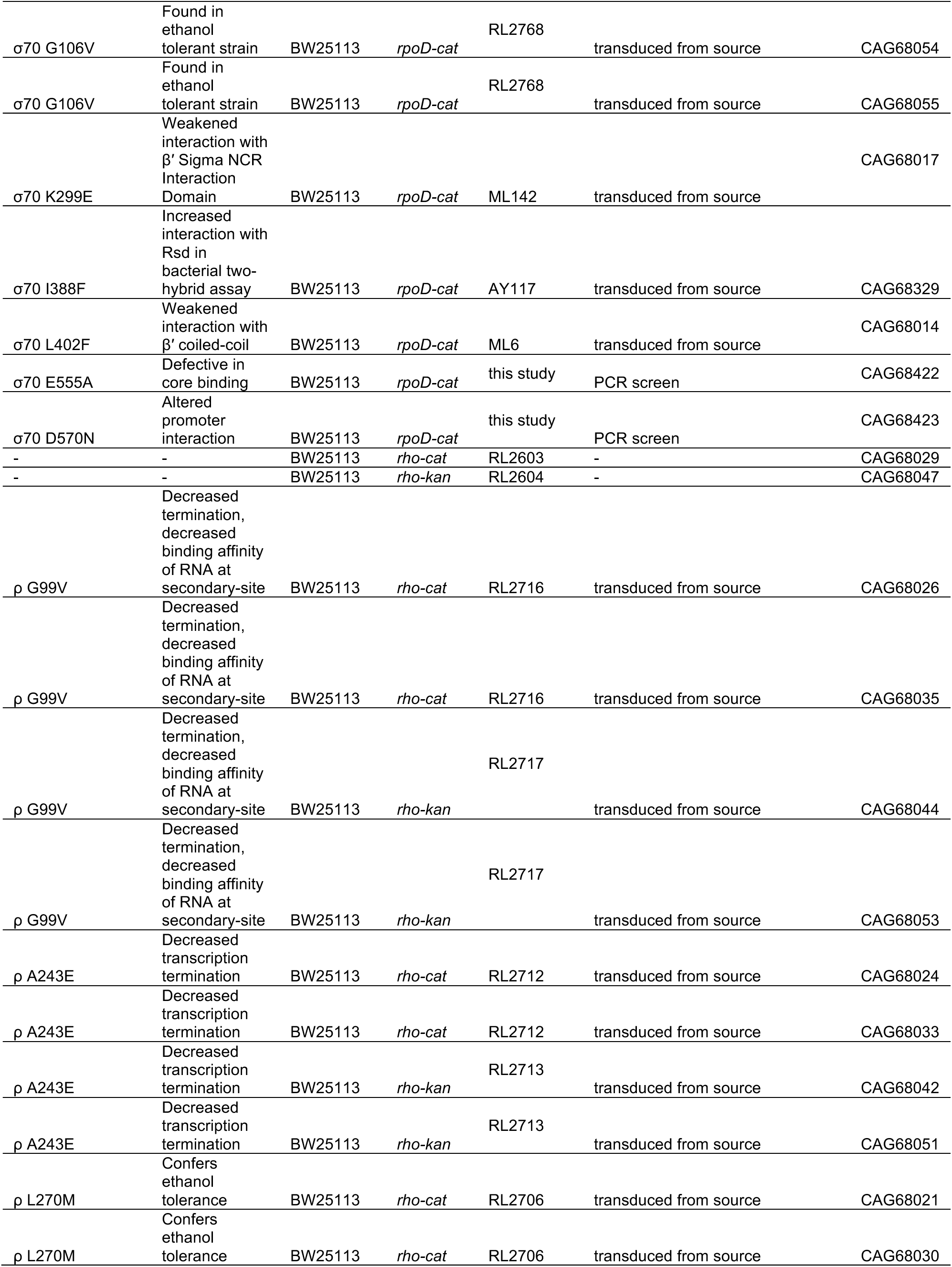
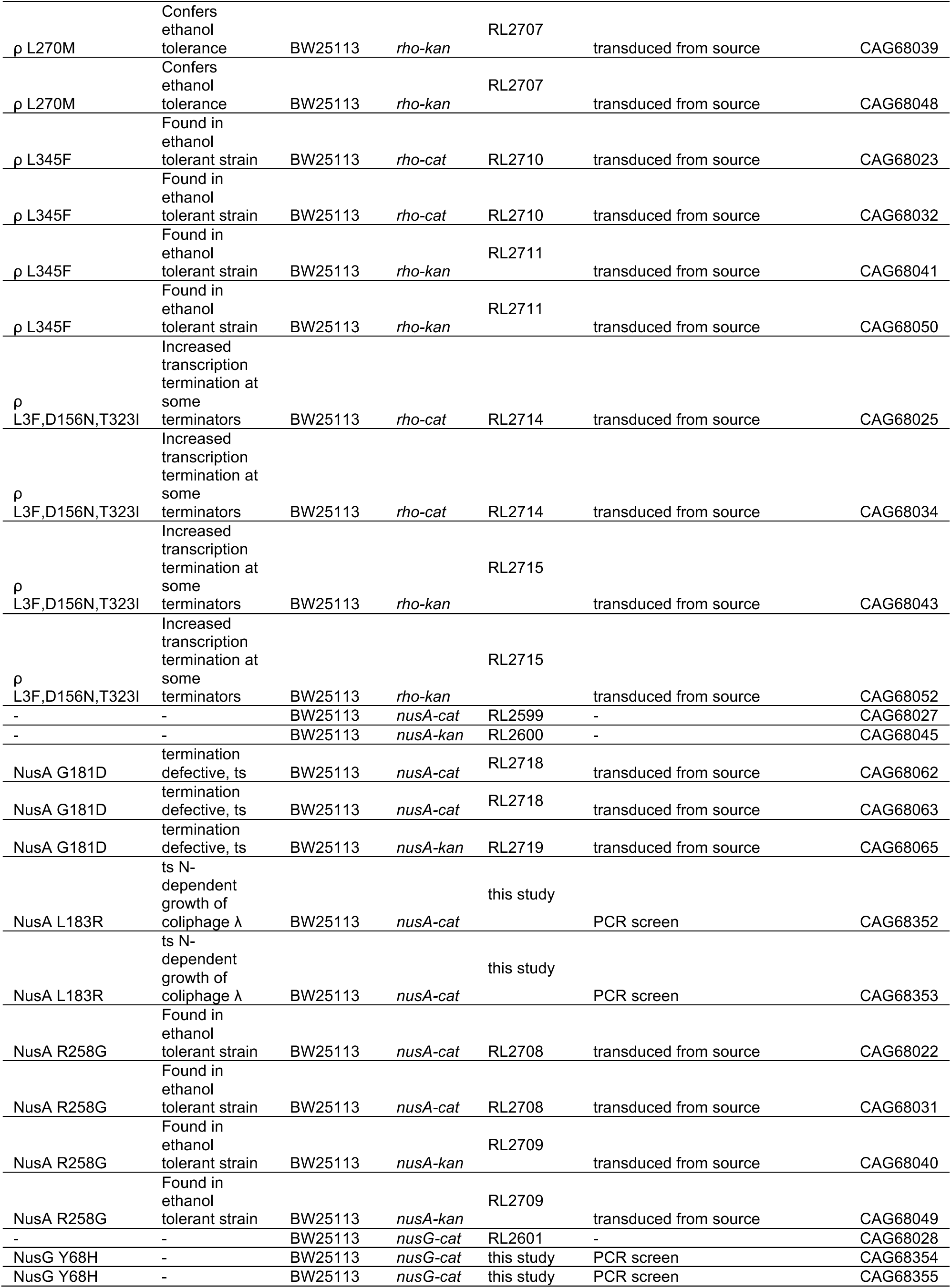
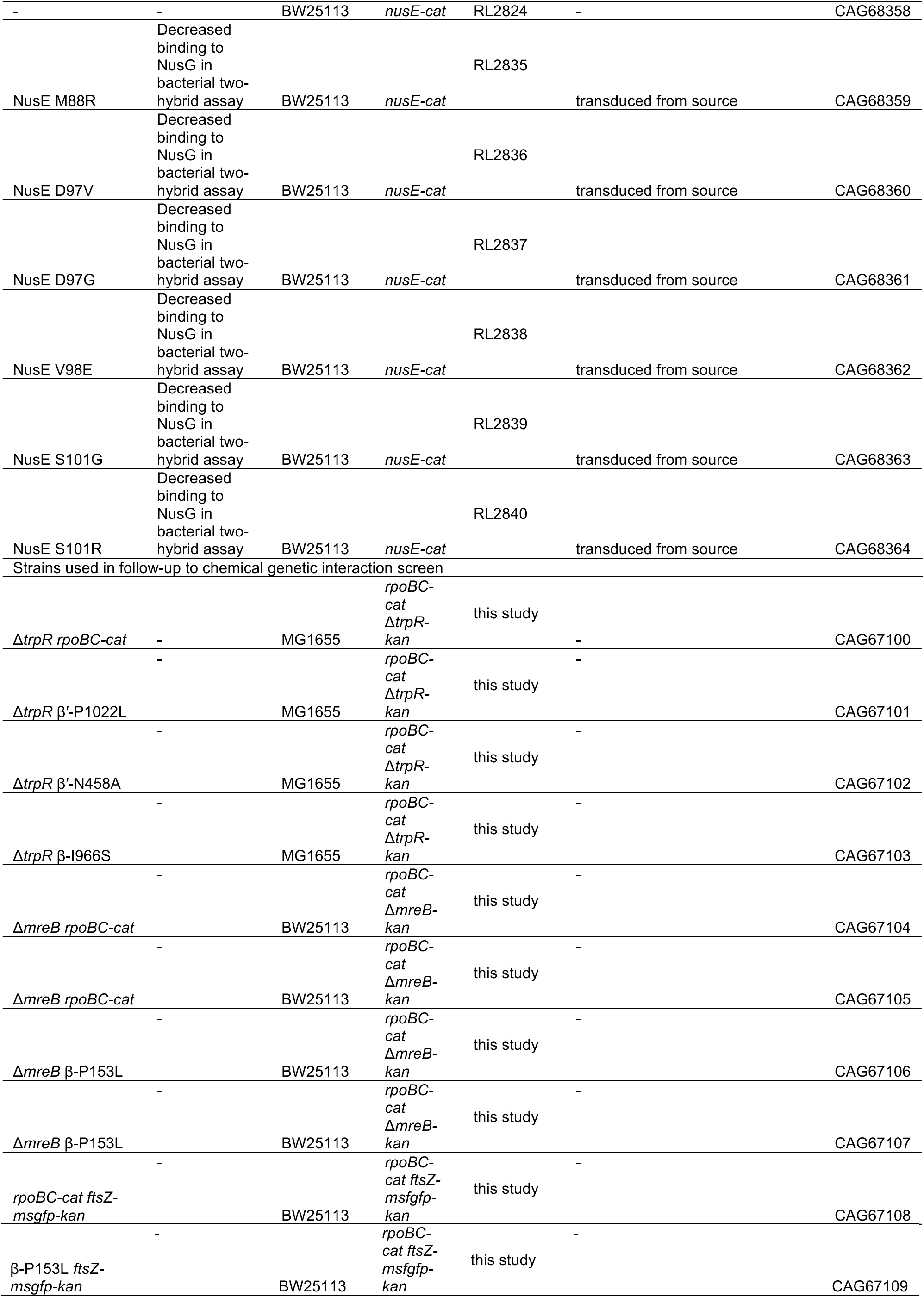
Strains used in this study

**Supplemental Table 2:** Conditional enrichments among transcription mutant clusters

| Condition | NES | p-adj | individual significant |
| --- | --- | --- | --- |
| **Cluster15** |  |  |  |
| isopentanol | -1.79 | $2.03 \times 10^{-2}$ | 1 |
| t-butanol | -1.7 | $3.04 \times 10^{-2}$ | 2 |
| azithromycin | 1.73 | $9.94 \times 10^{-3}$ | 1 |
| guanidine hydrochloride | 1.77 | $1.61 \times 10^{-2}$ | 2 |
| pseudomonic acid A | 1.82 | $9.38 \times 10^{-3}$ | 5 |
| tetracycline | 1.85 | $3.04 \times 10^{-2}$ | 1 |
| doxycycline | 1.99 | 0.00 | 4 |
| **Cluster12** |  |  |  |
| **Cluster13** |  |  |  |
| **Cluster16** |  |  |  |
| tetracycline | -1.97 | $1.00 \times 10^{-2}$ | 1 |
| pseudomonic acid A | -1.82 | $2.42 \times 10^{-2}$ | 3 |
| D,L-serine hydroxamate | -1.82 | $2.00 \times 10^{-2}$ | 2 |
| doxycycline | -1.77 | $2.73 \times 10^{-2}$ | 1 |
| sulfamonomethoxine | 2.08 | 0.00 | 1 |
| **Cluster14** |  |  |  |
| chlorhexidine dihydrochloride | -2.1 | 0.00 | 2 |
| azelaic acid | -2.02 | 0.00 | 8 |
| spectinomycin | -2.01 | 0.00 | 8 |
| gentamicin | -1.94 | 0.00 | 5 |
| tobramycin | -1.84 | $3.58 \times 10^{-3}$ | 7 |
| silver(II) | -1.82 | 0.00 | 3 |
| copper(II) | -1.76 | $2.03 \times 10^{-2}$ | 4 |
| holomycin | -1.75 | $3.04 \times 10^{-2}$ | 1 |
| polymyxin B | -1.74 | $1.00 \times 10^{-2}$ | 8 |
| isopropanol | -1.74 | $1.15 \times 10^{-2}$ | 6 |
| 25C | -1.71 | $1.11 \times 10^{-2}$ | 10 |
| M9min casamino acids | -1.7 | $1.88 \times 10^{-2}$ | 8 |
| tunicamycin | -1.68 | 0.00 | 7 |
| ethanol | -1.66 | $8.21 \times 10^{-3}$ | 11 |
| n-butanol | -1.61 | $2.00 \times 10^{-2}$ | 4 |
| M9min glycerol | 1.62 | $4.03 \times 10^{-2}$ | 3 |
| M9min maltose | 1.64 | $4.75 \times 10^{-2}$ | 7 |
| M9min glucosamine | 1.66 | $2.37 \times 10^{-2}$ | 6 |
| M9min succinate | 1.79 | $9.89 \times 10^{-3}$ | 6 |
| M9min 5-fluorouridine | 1.82 | $2.21 \times 10^{-2}$ | 2 |
| M9min 5-methyltryptophan | 1.82 | $9.94 \times 10^{-3}$ | 2 |
| phenazine methosulfate | 1.85 | $1.55 \times 10^{-2}$ | 3 |
| bleomycin | 1.95 | 0.00 | 1 |
| mecillinam | 1.96 | $3.97 \times 10^{-3}$ | 12 |
| A22 | 2.08 | 0.00 | 5 |
| **Cluster2** |  |  |  |
| **Cluster23** |  |  |  |
| rifampicin | -2.03 | 0.00 | 4 |
| mecillinam | -2 | $4.08 \times 10^{-2}$ | 1 |
| sodium fluoride | -1.91 | $2.73 \times 10^{-2}$ | 3 |
| ciprofloxacin | -1.86 | $1.00 \times 10^{-2}$ | 1 |
| ethidium bromide | -1.81 | $3.04 \times 10^{-2}$ | 2 |
| azidothymidine | -1.75 | $3.04 \times 10^{-2}$ | 2 |
| spiramycin | -1.72 | $3.40 \times 10^{-2}$ | 1 |
| dibucaine | -1.63 | $3.18 \times 10^{-2}$ | 1 |
| bicyclomycin | -1.6 | $1.25 \times 10^{-2}$ | 5 |
| cinoxacin | 1.66 | $3.95 \times 10^{-2}$ | 1 |
| **Cluster20** |  |  |  |
| rifampicin | 1.61 | $7.91 \times 10^{-3}$ | 4 |
| chlorhexidine dihydrochloride | 1.72 | $4.75 \times 10^{-2}$ | 1 |

**Supplemental Table 3:** Conditional enrichments among pre-defined transcription mutant classes

| Condition | NES | p-adj | individual significant |
| --- | --- | --- | --- |
| **M+** |  |  |  |
| t-butanol | -2.16 | 0.00 | 2 |
| holomycin | -2.14 | 0.00 | 2 |
| isopropanol | -2.08 | 0.00 | 8 |
| isopentanol | -2.02 | $4.10 \times 10^{-3}$ | 1 |
| polymyxin B | -2.01 | 0.00 | 9 |
| 25C | -1.88 | 0.00 | 13 |
| chlorhexidine dihydrochloride | -1.86 | $3.40 \times 10^{-2}$ | 2 |
| 10C | -1.8 | $6.04 \times 10^{-3}$ | 7 |
| n-butanol | -1.78 | 0.00 | 9 |
| M9min casamino acids | -1.75 | $1.25 \times 10^{-2}$ | 10 |
| azelaic acid | -1.73 | $1.90 \times 10^{-2}$ | 7 |
| tunicamycin | -1.72 | 0.00 | 8 |
| silver(II) | -1.71 | $2.03 \times 10^{-2}$ | 4 |
| spectinomycin | -1.63 | $3.04 \times 10^{-2}$ | 8 |
| ethanol | -1.56 | $2.87 \times 10^{-2}$ | 10 |
| M9min 5-methyltryptophan | 1.65 | $4.57 \times 10^{-2}$ | 4 |
| M9min glycerol | 1.68 | $2.03 \times 10^{-2}$ | 3 |
| bleomycin | 1.73 | $3.40 \times 10^{-2}$ | 1 |
| M9min 5-methylanthranilic acid | 1.77 | $1.90 \times 10^{-2}$ | 9 |
| mecillinam | 1.83 | $1.00 \times 10^{-2}$ | 10 |
| guanidine hydrochloride | 1.88 | $4.82 \times 10^{-3}$ | 2 |
| pseudomonic acid A | 1.89 | $2.03 \times 10^{-2}$ | 9 |
| M9min glucosamine | 1.91 | 0.00 | 7 |
| M9min succinate | 1.91 | 0.00 | 7 |
| M9min maltose | 1.92 | 0.00 | 8 |
| doxycycline | 1.99 | $5.49 \times 10^{-3}$ | 4 |
| tetracycline | 1.99 | $1.55 \times 10^{-2}$ | 1 |
| A22 | 2.06 | 0.00 | 7 |
| **5-MAA-R** |  |  |  |
| **Ethanol tolerant** |  |  |  |
| sodium fluoride | -1.96 | $1.52 \times 10^{-2}$ | 3 |
| sulfamonomethoxine | -1.9 | 0.00 | 1 |
| azidothymidine | -1.81 | $2.73 \times 10^{-2}$ | 1 |
| hydroxyurea | -1.6 | $2.42 \times 10^{-2}$ | 4 |
| urea | -1.58 | $4.57 \times 10^{-2}$ | 4 |
| trimethoprim | -1.57 | $3.02 \times 10^{-2}$ | 5 |
| **Galr** |  |  |  |
| **Mpa** |  |  |  |

**Condition:** the name of the stressor, **NES:** the normalized enrichment score as output by GSEA v.3.0. Larger magnitude scores indicate a higher enrichment. Negative NES values correspond to an enrichment for negative S-scores. Positive NES values correspond to an enrichment for positive S-scores. Enriched conditions are sorted according to NES. **p-adj:** The Benjamini-Hochberg FDR corrected p-values (corrected from the nominal p-values output by GSEA). **individual significant:** The number of
mutants within the group with an S-score that was individually significant for this condition. **M+**: The class of M+ mutants. **5-MAA-R**: The class of known hyper-attenuation mutants, as defined resistance to 5-methyl-anthranilic acid in a $\Delta trpR$ background. **Ethanol tolerant**: Mutations found in strains evolved for tolerance to ethanol. **Galr**: suppression of galactose sensitivity in *gal10 $\Delta$ 56* in RNA polymerase II from *Saccharomyces cerevisiae*. **Mpa**: mycophenolic acid sensitivity in RNA polymerase II from *Saccharomyces cerevisiae*.

**Supplemental table 4:** Mutations detected in whole genome resequencing of Δ*mreB* strains

| position | mutation | annotation | gene | description | Notes |
| --- | --- | --- | --- | --- | --- |
| CAG67202, *rpoBC-cat* Parental strain: BW25113 (CAG67001) |  |  |  |  |  |
| 1,484,419 | Δ9 bp | pseudogene (544-552/1001 nt) | *gapC* ← | pseudogene, GAP dehydrogenase; glyceraldehyde-3-phosphate dehydrogenase (second fragment) | SNP |
| 4,029,087 | A→T | noncoding (197/1542 nt) | *rrsA* → | 16S ribosomal RNA of *rrnA* operon | SNP |
| 4,175,201 | +T | coding (4029/4029 nt) | *rpoB* → | RNA polymerase, beta subunit | SNP associated with cat marker insertion |
| 4,175,845 | +A | coding (609/639 nt) | *cat* → |  | SNP associated with cat marker insertion |
| 4,175,857 | +T | coding (621/639 nt) | *cat* → |  | SNP associated with cat marker insertion |
| 4,175,978 | +A | intergenic (+103/-20) | *cat* → / → *rpoC* | –/RNA polymerase, beta prime subunit | SNP associated with cat marker insertion |
| CAG67104 *rpoBC-cat ΔmreB*, Parental strain: CAG67202 |  |  |  |  |  |
| 1,484,419 | Δ9 bp | pseudogene (544-552/1001 nt) | *gapC* ← | pseudogene, GAP dehydrogenase; glyceraldehyde-3-phosphate dehydrogenase (second fragment) | Parental SNP |
| 4,029,087 | A→T | noncoding (197/1542 nt) | *rrsA* → | 16S ribosomal RNA of *rrnA* operon | Parental SNP |
| 4,175,201 | +T | coding (4029/4029 nt) | *rpoB* → | RNA polymerase, beta subunit | Parental SNP |
| 4,175,845 | +A | coding (609/639 nt) | *cat* → |  | Parental SNP |
| 4,175,857 | +T | coding (621/639 nt) | *cat* → |  | Parental SNP |
| 4,175,978 | +A | intergenic (+103/-20) | *cat* → / → *rpoC* | –/RNA polymerase, beta prime subunit | Parental SNP |
| 3393996-3394395 | Δ400 | missing coverage | *mreB* | cell wall structural complex MreBCD, actin-like component MreB | *ΔmreB::kan* |
| CAG67105 *rpoBC-cat ΔmreB*, Parental strain: CAG67202 |  |  |  |  |  |
| 1,484,419 | Δ9 bp | pseudogene (544-552/1001 nt) | *gapC* ← | pseudogene, GAP dehydrogenase; glyceraldehyde-3-phosphate dehydrogenase (second fragment) | Parental SNP |
| 4,029,087 | A→T | noncoding (197/1542 nt) | *rrsA* → | 16S ribosomal RNA of *rrnA* operon | Parental SNP |
| 4,175,201 | +T | coding (4029/4029 nt) | *rpoB* → | RNA polymerase, beta subunit | Parental SNP |
| 4,175,845 | +A | coding (609/639 nt) | *cat* → |  | Parental SNP |
| 4,175,857 | +T | coding (621/639 nt) | *cat* → |  | Parental SNP |
| 4,175,978 | +A | intergenic (+103/-20) | *cat* → / → *rpoC* | –/RNA polymerase, beta prime subunit | Parental SNP |
| 3394017-3394392 | Δ376 | missing coverage | *mreB* | cell wall structural complex MreBCD, actin-like component MreB | Δ*mreB*::kan |
| CAG68095 β-P153L, Parental strain: CAG67202 |  |  |  |  |  |
| 1,484,419 | Δ9 bp | pseudogene (544-552/1001 nt) | *gapC* ← | pseudogene, GAP dehydrogenase; glyceraldehyde-3-phosphate dehydrogenase (second fragment) | Parental SNP |
| 4,029,087 | A→T | noncoding (197/1542 nt) | *rrsA* → | 16S ribosomal RNA of *rrnA* operon | Parental SNP |
| 4,171,630 | C→T | P153L (CCG→CTG) | *rpoB* → | RNA polymerase, beta subunit | β-P153L |
| 4,175,201 | +T | coding (4029/4029 nt) | *rpoB* → | RNA polymerase, beta subunit | Parental SNP |
| 4,175,845 | +A | coding (609/639 nt) | *cat* → |  | Parental SNP |
| 4,175,857 | +T | coding (621/639 nt) | *cat* → |  | Parental SNP |
| 4,175,978 | +A | intergenic (+103/-20) | *cat* → / → *rpoC* | –/RNA polymerase, beta prime subunit | Parental SNP |
| CAG667106 β-P153L Δ*mreB*, Parental strain: CAG68095 |  |  |  |  |  |
| 1,484,419 | Δ9 bp | pseudogene (544-552/1001 nt) | *gapC* ← | pseudogene, GAP dehydrogenase; glyceraldehyde-3-phosphate dehydrogenase (second fragment) | Parental SNP |
| 3,430,056 | G→A | E60E (GAG→GAA) | *trkA* → | NAD-binding component of Trk potassium transporter | SNP in BW2592 genetically linked to *mreB* |
| 4,029,087 | A→T | noncoding (197/1542 nt) | *rrsA* → | 16S ribosomal RNA of *rrnA* operon | parental SNP |
| 4,171,630 | C→T | P153L (CCG→CTG) | *rpoB* → | RNA polymerase, beta subunit | parental SNP |
| 4,175,201 | +T | coding (4029/4029 nt) | *rpoB* → | RNA polymerase, beta subunit | parental SNP |
| 4,175,845 | +A | coding (609/639 nt) | *cat* → |  | parental SNP |
| 4,175,857 | +T | coding (621/639 nt) | *cat* → |  | parental SNP |
| 4,175,978 | +A | intergenic (+103/-20) | *cat* → / → *rpoC* | –/RNA polymerase, beta prime subunit | parental SNP |
| 3393999-3394409 | Δ411 | missing coverage | *mreB* | cell wall structural complex MreBCD, actin-like component MreB | Δ*mreB*::kan |
| CAG67107 β-P153L Δ*mreB*, Parental strain: CAG68095 |  |  |  |  |  |
| 1,484,419 | Δ9 bp | pseudogene (544-552/1001 nt) | *gapC* ← | pseudogene, GAP dehydrogenase; glyceraldehyde-3-phosphate dehydrogenase (second fragment) | Parental SNP |
| 3,430,056 | G→A | E60E (GAG→GAA) | *trkA* → | NAD-binding component of Trk potassium transporter | SNP in BW2592 genetically linked to *mreB* |
| 3,467,784 | T→C | K43R (AAA→AGA) | *rpsL* ← | 30S ribosomal subunit protein S12 | SNP in BW2592 genetically linked to *mreB* |
| 4,029,087 | A→T | noncoding (197/1542 nt) | *rrsA* → | 16S ribosomal RNA of *rrnA* operon | parental SNP |
| 4,171,630 | C→T | P153L (CCG→CTG) | *rpoB* → | RNA polymerase, beta subunit | parental SNP |
| 4,175,201 | +T | coding (4029/4029 nt) | *rpoB* → | RNA polymerase, beta subunit | parental SNP |
| 4,175,845 | +A | coding (609/639 nt) | *cat* → |  | parental SNP |
| 4,175,857 | +T | coding (621/639 nt) | *cat* → |  | parental SNP |
| 4,175,978 | +A | intergenic (+103/-20) | *cat* → / → *rpoC* | –/RNA polymerase, beta prime subunit | parental SNP |
| 3393993-3394395 | Δ403 | missing coverage | *mreB* | cell wall structural complex MreBCD, actin-like component MreB | Δ*mreB*::kan |

**Position:** Position in the genome of the detected mutation. **Mutation:** The identity of the detected mutation **Annotation:** The class of mutation detected (amino acid change, coding, intergenic, and missing coverage) **Gene:** The gene affected by the mutation (or those nearby if intergenic) **Description:** Description of the affected gene. **Notes:** Annotation for the source of the detected SNP (parental SNP, derived from the BW2592 donor strain, or a true mutation)

**Supplemental Table 5:**
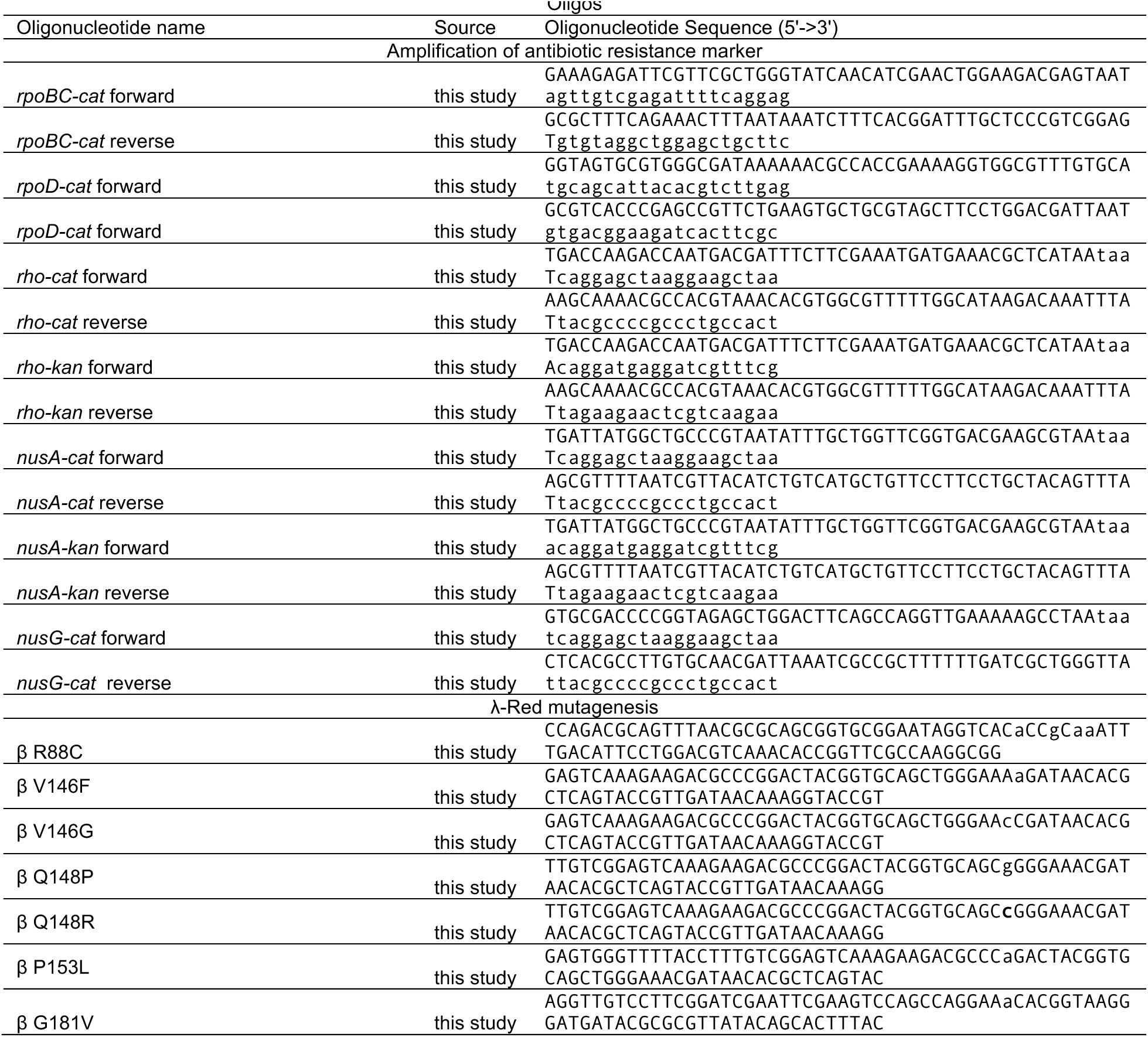
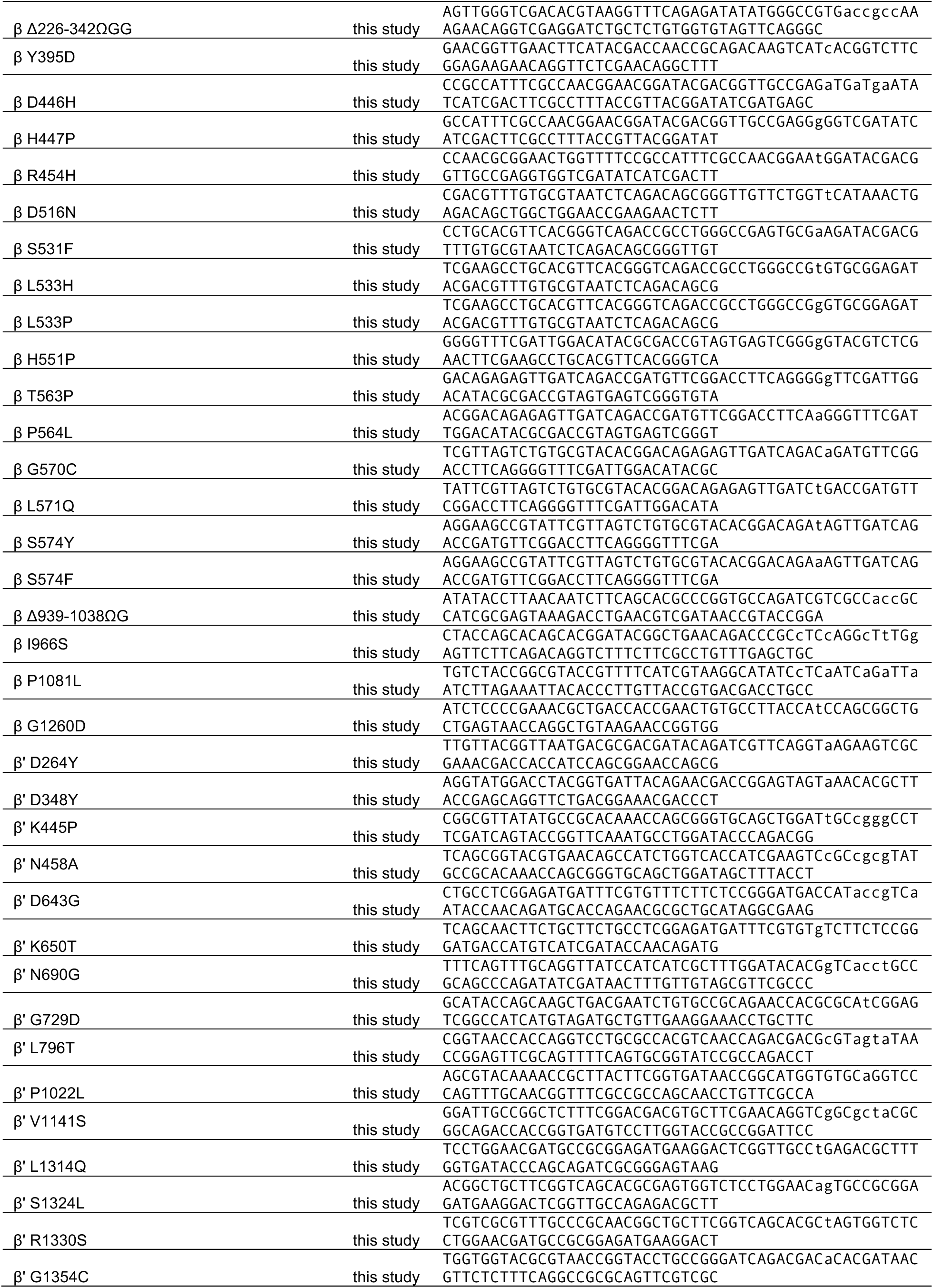
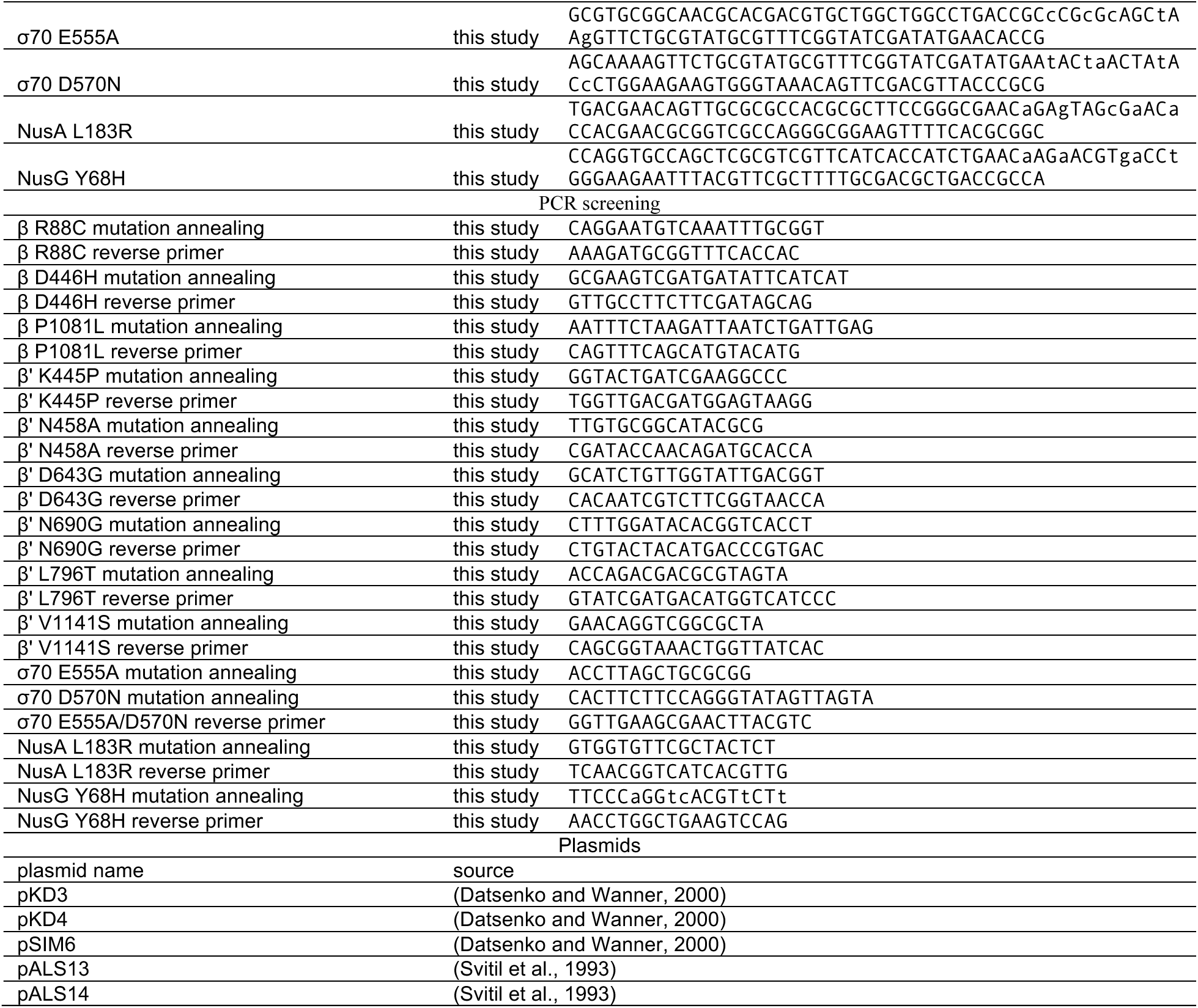
Oligos and plasmids used in this study

